# VPRBP functions downstream of the androgen receptor and OGT to restrict p53 activation in prostate cancer

**DOI:** 10.1101/2021.02.28.433236

**Authors:** Ninu Poulose, Nicholas Forsythe, Adam Polonski, Gemma Gregg, Sarah Maguire, Marc Fuchs, Sarah Minner, Guido Sauter, Simon S. McDade, Ian G. Mills

## Abstract

A comprehensive understanding of androgen receptor (AR) signalling mechanisms during prostate carcinogenesis is instrumental in developing novel therapies. Studies have shown glycosylation as a key androgen regulated process and that O-GlcNAc transferase (OGT), the enzyme that catalyses the covalent addition of UDP-N-acetylglucosamine (UDP-GlcNAc) to serine and threonine residues of proteins, is often up-regulated in prostate cancer (PCa) with its expression correlated with high Gleason score. In this study we have identified an AR and OGT co-regulated factor, VPRBP also known as DCAF1. We show that VPRBP is regulated by the AR at the transcript level, and stabilized by OGT at the protein level. VPRBP knockdown in PCa cells led to a significant decrease in cell proliferation, p53 stabilization, nucleolar fragmentation and increased p53 recruitment to the chromatin. In human prostate tumor samples, VPRBP protein overexpression correlated with AR amplification, OGT overexpression, early biochemical relapse and poor clinical outcome. Analysis of TCGA datasets found a positive correlation of VPRBP with AR mRNA expression. Furthermore, AR activity gene signature analysis revealed a positive correlation of VPRBP with a subset of AR target genes implying that VPRBP has a preferential regulatory impact on part of the AR regulome. In conclusion, we have shown that VPRBP/DCAF1 promotes PCa cell proliferation by restraining p53 activation under the influence of the AR and OGT as well as uncovered a unique subset of AR activity gene signature that correlates with VPRBP expression with the potential for new avenues for patient stratification and treatment.

## Introduction

AR activity plays an important role in the development of localized prostate cancer and also in the sustaining treatment-resistant metastatic disease ^1^. AR activation has been shown to enhance flux through hexosamine biosynthetic pathway (HBP) in PCa cell lines ^2^, which leads to increased bioavailability of UDP-N-acetylglucosamine, a substrate for O-GlcNAcylation as well as N-linked and O-linked glycosylation ^3^. O-GlcNAcylation, a highly dynamic and often transient post-translational modification (PTM) is specifically increased in PCa tissues compared to adjacent non-malignant tissues ^4^. This PTM is regulated by two enzymes, OGT that catalyses the covalent addition of UDP-N-acetylglucosamine to serine and threonine residues of cytoplasmic, nuclear and mitochondrial proteins, and O-GlcNAcase (OGA) which removes the O-GlcNAc moiety ^5^. OGT is considered to be a metabolic rheostat and its expression is elevated in many cancers including PCa, with higher O-GlcNAc levels associated with poor prognosis of patients ^6 7^. There is a growing diverse list of proteins which undergo this PTM, including some of the key transcription factors such as c-Myc ^8^ and p53 ^9^.

A recent ChIP-seq study by Itkonen *et al* has demonstrated that the O-GlcNAc chromatin mark is rapidly lost upon inhibition of OGT activity by a fast-acting inhibitor, OSMI2 ^10^ in PCa cells. This analysis revealed that the majority of the O-GlcNAc peaks were promoter associated with over 95% overlap with DNase-hypersensitive regions and active chromatin marks. Independent AR ChIP-seq studies from our lab and others have shown that the majority of AR binding sites are distal intergenic and intronic ^11^. Genome-wide motif co-enrichment, have shown entirely distinct associations between O-GlcNAc and other factors (principally c-Myc and ETS transcription factors), and the AR and other factors (Forkhead family transcription factors such as FOXA1)^11^. Despite these differences, we know that both AR and OGT contribute to PCa progression. To better understand the interplay between OGT and AR in PCa, we analysed these AR and O-GlcNAc ChIP-seq data focusing on promoter proximal sites and identified a small number of overlapping sites and genes. We focussed on VPRBP (Vpr binding protein) also known as DCAF1 (DDB1 and CUL4 Associated Factor 1) which has been implicated as a regulator of cell cycle and cell proliferation^12,13^. VPRBP is the substrate recognition component of cullin 4A-ring E3 ubiquitin ligase (CRL4A) complex as well as separate HECT type EDD/UBR5 E3 ligase ^14^. In this study we show that VPRBP is a novel AR target as well as an OGT regulated protein. Knockdown of VPRBP led to a marked reduction in PCa cell proliferation. We go on to show that VPRBP down-regulates p53 stability and activity, and that this is in part by maintaining nucleolar integrity. Tissue microarray studies showed a positive correlation of VPRBP expression with AR/OGT expression and an inverse correlation with PSA recurrence free survival. Furthermore, VPRBP expression in TCGA datasets showed a positive correlation with AR expression and a subset of AR activity gene signatures, and inverse correlation with the p53 pathway. We conclude that VPRBP acts as a novel downstream effector of AR and OGT mediated PCa cell proliferation by impairing p53 checkpoint activation.

## Results

### Identification of VPRBP as a novel AR regulated gene

To identify AR and OGT co-regulated genes, we analysed published AR (GSE28126) and O-GlcNAc (GSE112667) ChIP-seq data. Comparing peak distribution between AR and O-GlcNAc binding sites from these two separate studies (Fig. 1A), indicated that, as previously reported the majority of O-GlcNAc binding sites are promoter proximal, whereas the majority of AR binding sites are intronic or associated with distal inter-genic regions (Fig. 1A). Intersecting LNCaP AR ChIP-seq (R1881 stimulated) binding sites with O-GlcNAc ChIP-seq consensus sites identified only 9 overlapping sites, amongst which a binding site was detected within proximity of *VPRBP* gene (Fig. 1B). VPRBP is of particular interest because it has been shown to be highly expressed in different tumor tissues ^15^ and is also known to play a pivotal role in cell growth and cell cycle entry in T cells ^12^. Moreover, depletion of VPRBP in DU145 prostate cancer cells reduced cell proliferation and number of colony forming cells ^15^. However the roles of VPRBP in mediating androgen response in PCa or its regulation by OGT or O-GlcNAcylation have not been reported so far.

**Figure 1.**
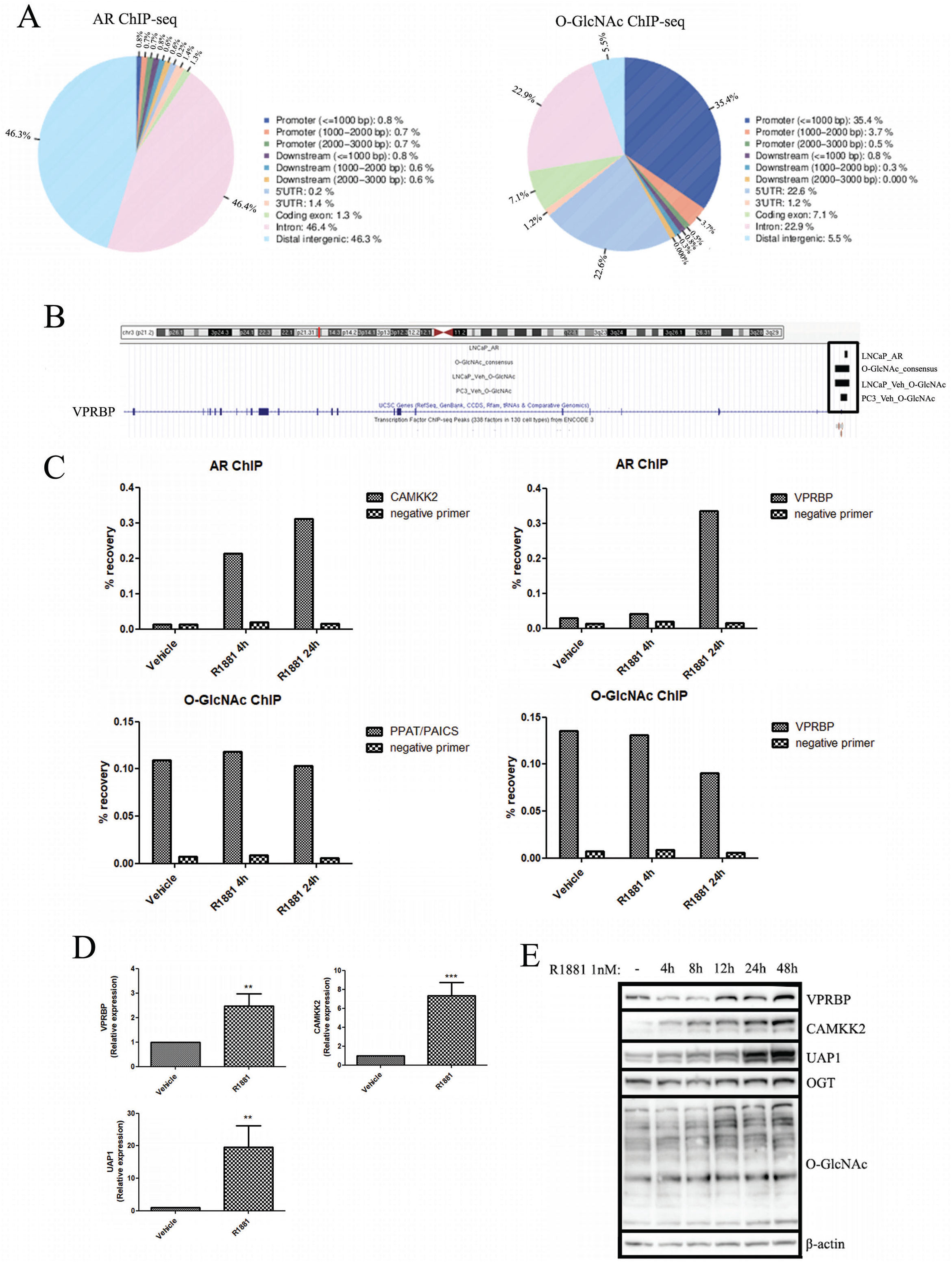
Identification of VPRBP as an AR and O-GlcNAc co-regulated target. (A) Venn diagrams showing the distribution of peaks in relation to genes in AR and O-GlcNAc consensus ChIP-seq, generated using CEAS tool in galaxy cistrome. (B) ChIP-seq enrichment of AR and O-GlcNAc at VPRBP promoter region using UCSC genome browser. (C) Percentage recovery of CAMKK2 and VPRBP with AR ChIP; and PPAT/PAICS and VPRBP with O-GlcNAc ChIP in vehicle (0.01% ethanol) and 1nM R1881 4h and 24h treated LNCaP cells. (D) mRNA expression of VPRBP, CAMKK2 and UAP1 in LNCaP treated with vehicle or 1nM R1881 for 24 h was detected by qRT-PCR. Results are normalized to RPLPO as housekeeping control. (E) Time dependency of VPRBP expression following exposure to 1nM R1881 for different time points. LNCaP cells were androgen deprived for 3 days prior to stimulation with vehicle or 1 nM R1881. p values by Student’s t test. ** =p < 0.01; *** = p < 0.001.

To confirm AR and O-GlcNAc enrichment at the VPRBP site, we performed ChIP-qPCR in LNCaP cells stimulated with 1nM R1881, a synthetic androgen, for 4h and 24h following 72h of androgen deprivation ^11111^. We confirmed that as expected androgen stimulation resulted in AR enrichment at a CAMKK2 associated site (a known AR target ^11^), at both time points (Fig. 1C). By contrast O-GlcNAc enrichment in the promoter of the *PPAT/PAICS* gene ^16^ was not significantly altered in response to androgen treatment (Fig. 1C and Fig. S1A). Androgen stimulation resulted in increased binding of AR at the VPRBP promoter region at 24h time point (Fig. 1C and Fig. S1A).

Quantitative RT-PCR(qRT-PCR) analysis revealed that R1881 stimulated binding of AR to VPRBP promoter correlated with a 2.5 fold increase in VPRBP mRNA expression at 24 h (Fig. 1D), concomitant with a significant increase in VPRBP protein levels 24 and 48 hours (Fig. 1E and Fig. S1B). OGT and O-GlcNAcylation levels also showed significant increases following R1881 stimulation at the later time points (Fig. 1E and Fig.S1B). CAMKK2 and UAP1 expression levels were used as positive controls ^11^. Since R1881 is a synthetic androgen, we also tested the effect of endogenous androgen, dihydrotesteosterone (DHT). Similarly to R1881, stimulation of LNCaP cells by DHT for 24h also increased VPRBP mRNA and protein expression (Fig. S1C and Fig. S1D).

### OGT is required for VPRBP stability

To determine if OGT is required for VPRBP expression, we performed OGT knockdown using siRNA. This revealed that there was no significant change in basal VPRBP mRNA levels in OGT siRNA transfected cells compared to scrambled control (Fig. 2A). Interestingly, OGT knockdown reduced both basal (Fig. 2B) and androgen-induced (Fig. 2C) VPRBP expression. OGT siRNA transfection resulted in >90% reduction in basal OGT protein expression and >80% reduction in total O-GlcNAc levels with both siRNAs (Fig. S2A). There was ~57% reduction in VPRBP protein expression with OGTsiRNA1 and ~54% with OGT siRNA2 5d post transfection (Fig. S2A). Overall, these results suggest a possible dual regulation of VPRBP, whereby VPRBP is induced at the mRNA level by AR and is stabilised post-translationally by OGT activity.

**Figure 2.**
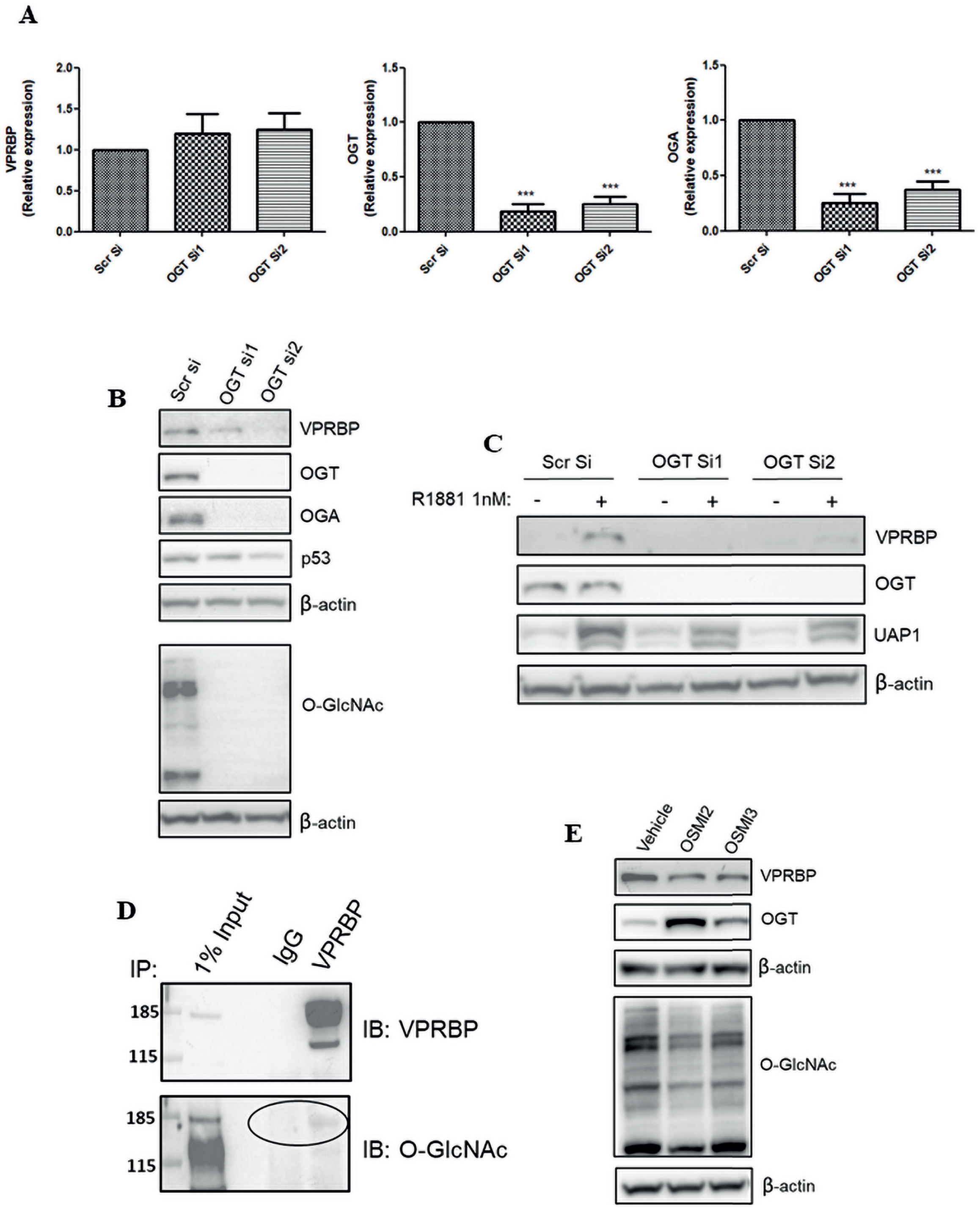
OGT is required for VPRBP stability. (A) LNCaP cells were transiently transfected with OGT siRNA and cells were harvested 5d post transfection to detect mRNA expression of VPRBP, OGT and OGA by qRT-PCR and (B) protein expression by immunoblot analysis under basal conditions. (C) For detection of protein expression under androgen stimulated conditions, LNCaP cells were transfected with OGT siRNAs or scrambled siRNA (scr si) followed by androgen deprivation for 72 h prior to 1nM R1881 stimulation for 24h. (D) The O-GlcNAcylation of immunoprecipitated (IP) VPRBP from LNCaP cells was detected by immunoblotting (IB) with RL2 antibody. (E) The effect of OGT inhibitors 40 μM OSMI2 and 10 μM OSMI3 on VPRBP protein levels following 24h treatment was detected by immnuoblot analysis. Results are expressed as means ± SD from at least three independent experiments. *** = p < 0.001 by Student’s t test.

O-GlcNAcylation has been shown to affect the stability of proteins like p53 ^9^, c-Myc ^2^ and EZH2 ^17^. Therefore we performed immunoprecipitation (IP) to test the O-GlcNAcylation status of VPRBP. IP indicates that VPRBP is an O-GlcNAcylated protein (Fig. 2D). To further confirm if O-GlcNAcylation is necessary for the stability of VPRBP, we treated the cells with inhibitors of OGT activity, OSMI2 and OSMI3 ^18^. We found that treatment for 24h decreased VPRBP expression at the protein level by ~73% for 40 μM OSMI2 and ~62% for 10 μM OSMI3 (Fig. 2E and Fig. S2B). OSMI treatment reduced overall O-GlcNAcylation levels with a compensatory up-regulation of OGT expression (Fig. 2E and Fig. SB). There was no significant change in VPRBP expression at the transcript level with OSMI3 treatment, although OSMI2 exhibited a small reduction of VPRBP transcripts (Fig. S2C).

### VPRBP down-regulation stabilizes p53 and inhibits LNCaP cell proliferation

Having identified VPRBP as an AR and OGT target, we went on to determine the functional effect of VPRBP knockdown in LNCaP cells. Knockdown with VPRBP siRNA resulted in ~75% reduction in VPRBP mRNA expression (Fig. 3A). A ~57% reduction in cell number was observed with VPRBP siRNAs when cells were grown in complete growth media containing androgens whereas androgen deprivation on its own resulted ~38% in a reduction in cell numbers (Fig. 3B). Furthermore, chemical inhibition by B32B3, a potent and selective inhibitor of VPRBP kinase activity ^15^, led to ~64% decrease in LNCaP cell proliferation at 5μM concentration (Fig. S3A). The combination of B32B3 and OGT inhibitors did not show significant decreases in cell numbers compared to B32B3 alone (Fig. S3A).

**Figure 3.**
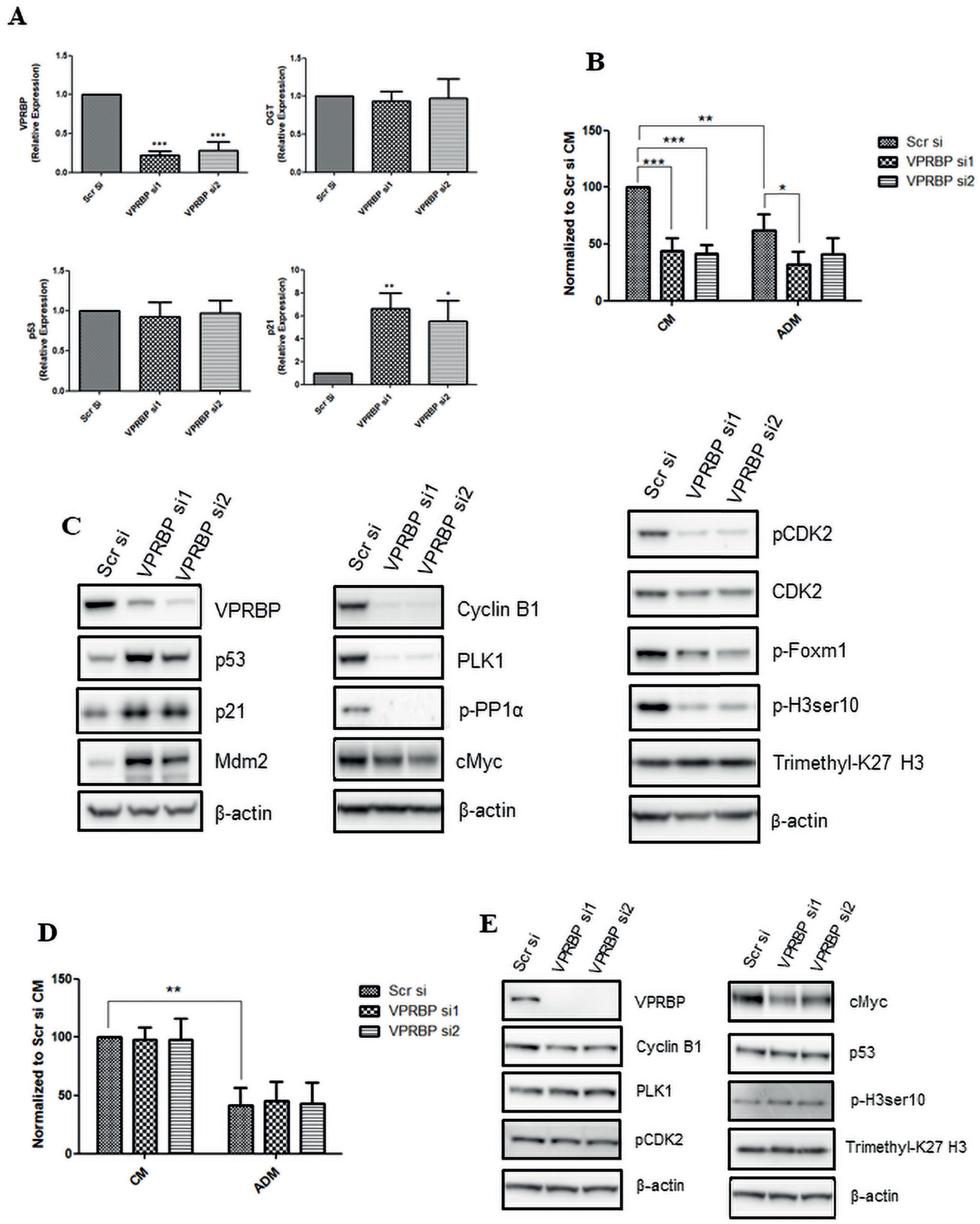
VPRBP knockdown leads to reduced cell proliferation and p53 stabilization. (A) LNCaP cells were transfected with VPRBP siRNA and cells were harvested 3d post transfection to detect mRNA expression of VPRBP, OGT, p53 and p21. (B) Effect of VPRBP knockdown on LNCaP cell proliferation was assessed by cell counting 5d post transfection; the cells were grown in the presence (CM) and absence of androgens (ADM). (C) Effect of VPRBP knockdown on LNCaP p53, markers of cell cycle and other proteins of interest was assessed by immunoblot analysis of cell lysate prepared 3d post transfection. (D) Effect of VPRBP knockdown on cell proliferation was assessed by cell counting 5d post transfection in VCaP cells grown in the presence (CM) and absence of androgens (ADM). (E) Effect of VPRBP knockdown on VCaP proteins of interest was assessed by immunoblot analysis. Results are expressed as means ± SD. * =p < 0.05, ** =p < 0.01; *** = p < 0.00. Statistical analyses were performed by Student’s t test for qPCR and one-way ANOVA followed by Tukey’s posthoc analysis for cell proliferation.

VPRBP knockdown in LNCaP led to a substantial increase in p53 protein expression and its downstream targets cyclin dependent kinase inhibitor p21^19^ and Mdm2^20^ (Mouse double minute 2 homolog), (Fig. 3C and Fig. S3B). Consequently we observed a drastic reduction in cell cycle markers like phospho CDK2 Thr160 ^21^, Cyclin B1 ^22^, polo-like kinase1 (PLK1) ^23^, phospho-histone H3 ser10 ^24^ and Phospho-PP1α Thr320 ^25^ (Fig. 3C and Fig. S3B) following VPRBP knockdown. In summary, VPRBP tightly control cell proliferation mainly by regulating the expression of p53. It is interesting to note that whereas VPRBP knockdown led to p53 stabilization, the reduction in VPRBP levels by OGT knockdown did not translate to similar effects (Fig. 2B). This is likely because p53 itself is an O-GlcNAcylated protein whose stability is enhanced with O-GlcNAcylation at Ser 149 position ^9^.

In order to determine if the growth inhibitory effects of VPRBP knockdown were primarily mediated through p53, we performed VPRBP knockdown in the VCaP, a cell-line expressing mutant p53. VPRBP knockdown in VCaP failed to cause significant changes in cell numbers (Fig. 3D) and cell cycle markers (Fig. 3E), indicating a crucial role for p53 in mediating VPRBP effects in PCa. However, similarly to LNCaP, VPRBP knockdown in VCaP cells reduced c-Myc levels (Fig. 3E). Together, these results suggest that the growth inhibitory effects of VPRBP knockdown are primarily mediated through p53 stabilization and activation.

Furthermore, we observed a reduction in p53 protein (Fig. S4A) and mRNA expression (Fig. S4B) in LNCaP cells following R1881 stimulation, and a corresponding decrease in p53 enrichment at p21 (CDKN1A) promoter by ChIP-qPCR (Fig. S4C) suggesting that pro-proliferative effects of androgens may partly be mediated by factors that down-regulate p53. However VPRBP gene itself does not appear to be p53 regulated, since there was no significant enrichment by ChIP-qPCR (Fig. S4D) and no binding sites were identified within its promoter region from p53 ChIP-seq in nutlin-3a treated LNCaP cells (Fig. S4E). Analysis of a novel p53-knockout LNCaP cell-line showed higher basal levels of VPRBP. By contrast treating LNCaP cells with a p53 stabilizer, nutlin-3a, led to a decrease in VPRBP (Fig. S4F and Fig. S4G). Consequently a reciprocal feedback relationship between VPRBP and p53 expression seems to exist within these cells.

### p53 ChIP-seq reveals increased p53 recruitment to the chromatin following VPRBP knockdown

To further establish whether VPRBP was influencing p53 activity, we performed p53 ChIP-seq in LNCaP cells following siRNA knockdown with VPRBP, OGT and non-targeting siRNAs along with nutlin-3a treated samples. Canonical p53 target gene p21 (CDKN1A) was used as a positive control to validate ChIP efficiency by ChIP-qPCR (Fig. 4A). This is the first report on p53 ChIP-seq in a prostate cancer cell line. Overall, there were fewer p53 peaks in LNCaP (Fig. 4B) than reported in other cell types ^26,27^, suggesting fundamental differences in p53 transcriptional landscape amongst these cells. ChIP-seq in nutlin-3a treated LNCaP cells returned 582 peaks, the majority of which (>80%) overlapped with nutlin-3a p53 ChIP-seq binding sites in other cell lines confirming they are bona fide p53 binding sites (Fig. 4C and Fig. S5A). By contrast VPRBP knockdown led to a ~4.7 fold increase in the number of p53 genomic binding sites compared to the scrambled control (Fig. 4B). There were 1387 consensus p53 binding sites between the two VPRBP knockdown samples (si1 and si2) representing a site overlap between these conditions of ~85%. The vast majority (~95%) of LNCaP nutlin-3a p53 ChIP-seq sites were present within this consensus suggesting that VPRBP knockdown expanded the number of p53 accessible sites (Fig. 4D). Importantly the majority of sites in VPRBP knockdown overlapped with nutlin-3a MCF7 p53 ChIP-seq peaks (Fig. 4E). The majority of OGT si1 and OGT si2 sites overlapped, so was there a significant overlap of OGT si consensus sites with VPRBP si consensus sites and scr si sites (Fig. S5B).

**Figure 4.**
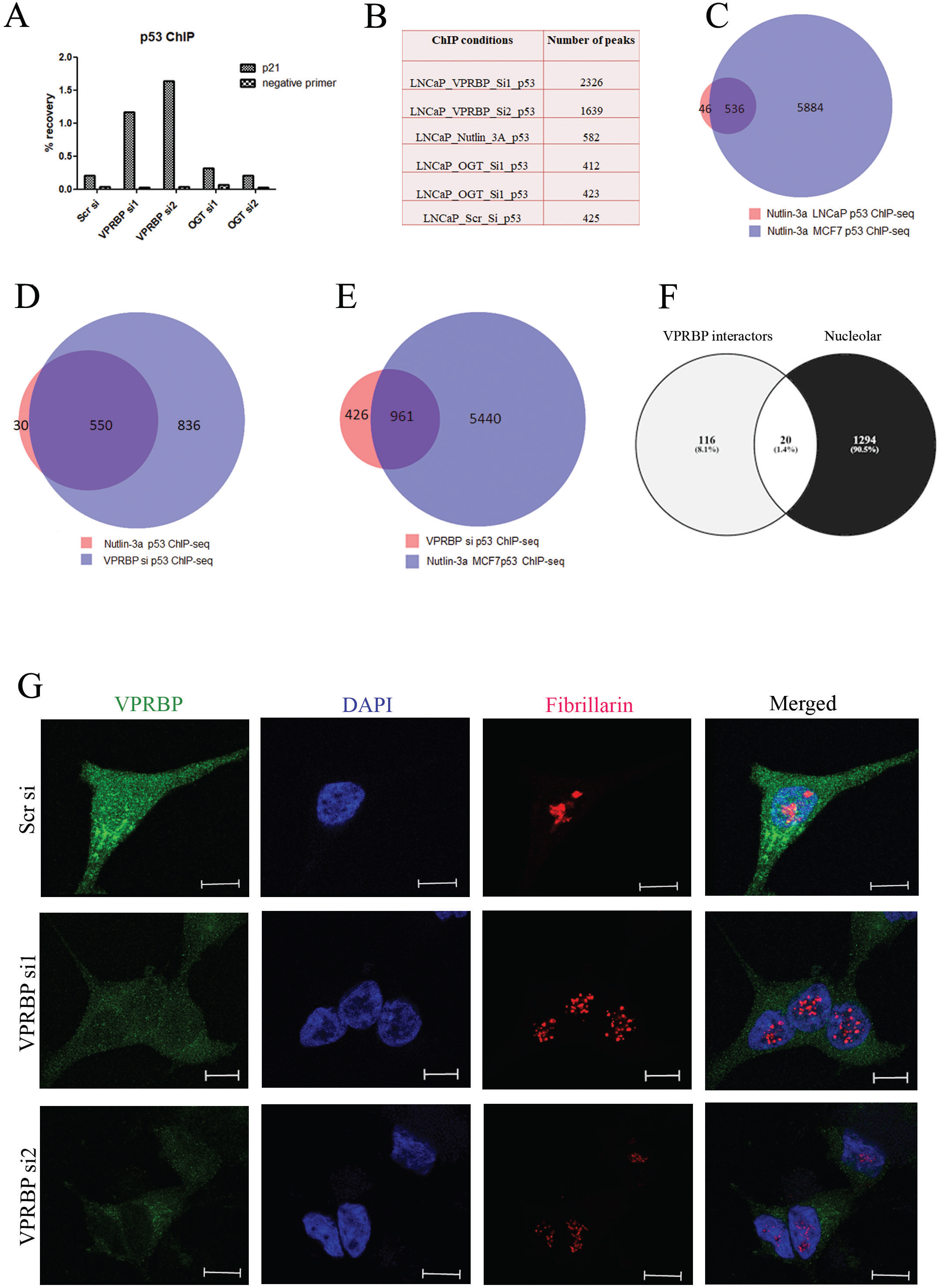
VPRBP knockdown increases p53 chromatin recruitment and induces nucleolar stress. (A) Bar graph showing percentage recovery of p21 and negative site primer (CCND1) following p53 ChIP in different transfection conditions. (B) Table showing the number of peaks obtained under different conditions. (C) Venn diagram showing the overlap of our nutlin-3a p53 ChIP-seq in LNCaP cells with previously reported nutlin-3a p53 ChIP-seq in MCF7 cells (GSE86164). (D) Venn diagram showing the overlap of nutlin-3a p53 ChIP-seq in LNCaP with VPRBPsi p53 ChIP-seq consensus sites. (E) Venn diagram showing the overlap of VPRBPsi p53 ChIP-seq consensus sites with previously reported nutlin-3a p53 ChIP-seq in MCF7 cells. (F) Venn diagram showing the overlap of VPRBP interactome with nucleolar proteome. (G) Representative immunofluorescence images showing VPRBP and fibrillarin staining in scrambled and VPRBP siRNA transfected LNCaPs 3d post transfection (scale bar= 10μm).

Assessing the distribution of p53 peaks revealed that the majority were present in distal intergenic regions (45-54%), with around 6% binding events in proximal promoters (Fig. S5C), resembling the AR binding pattern. The VPRBP si p53 ChIP-seq peaks showed ~53% overlap with H2K27Ac peaks and ~0.7% overlap with H3K27me3 peaks indicating majority of the binding sites are present at sites of transcriptional activity which are not accessible to p53 when VPRBP is expressed (Fig. S5D). The genes within 10kb of p53 sites in the VPRBP knockdown condition were identified (Supplementary file 2). These included established p53 target genes such as CDKN1A and MDM2. Motif enrichment analysis on the p53 sites in the knockdown condition showed an over-representation of TP53, TP73 and TP63 binding motifs (Fig. S5E). Together these suggest that VPRBP may restrain p53 activity by occluding chromatin and that VPRBP knockdown expands the gene regulatory network affected by increased p53 expression.

### VPRBP knockdown induces nucleolar stress in LNCaP cells

A comparison of VPRBP interactome (147 proteins) obtained from BioGRID database 28 (supplementary file 3) with nucleolar proteome (1314 proteins) of human cells derived from the Cell Atlas ^29^ showed that ~14 % of VPRBP interactors also show nucleolar localization (Fig. 4F). VPRBP is also reported to be involved in 40S ribosomal subunit biogenesis along with other CRL4 E3 ubiquitin ligase and COP9 signalosome components in a genome wide RNAi screen study ^30^. Hence we hypothesised that VPRBP knockdown may enhance p53 stability and diminish ribosome biogenesis by destabilising the nucleolus. Our immunofluorescence studies showed both cytoplasmic and nucleolar localization of VPRBP (Fig. 4G) as reported previously ^31 32^. Reduction in the number of nucleoli and/or disintegration of nucleolar structures are some of the characteristic features of nucleolar stress ^33^. We found a marked change in staining pattern of nucleolar protein fibrillarin following VPRBP siRNA knockdown, indicative of nucleolar stress (Fig. 4G). Fibrillarin showed exclusive nucleolar localization with a redistribution to small nucleoplasmic entities in the VPRBP knockdown cells, similar to previously described with actinomycin D treatment ^34^. A significant (~25%) reduction in fibrillarin and 40S ribosomal protein S8 (RPS8) expression, was observed with VPRBP siRNA2 (Fig. S6A and Fig. S6B.) This suggests that VPRBP knockdown contributes to p53 stabilization by imposing nucleolar stress.

### VPRBP expression correlates with AR expression in clinical samples and is prognostic

To further understand the prognostic potential of VPRBP in PCa, we analysed its expression in tissue microarrays (TMA) by IHC. The representative images of negative, low, intermediate and high staining of VPRBP are shown in Fig. S6C. We assessed its correlation with PCa pathology, and with the AR and OGT. We observed that 46% of all tumors stained strongly for VPRBP expression with a statistically significant increase in protein levels with tumor stage and quantitative Gleason grade (Supplementary Table 6). Although there were few Gleason score 7 tumors with tertiary Gleason pattern 5 (a powerful predictor of biochemical relapse ^35^), all of them were VPRBP-positive. A higher VPRBP staining was observed in cases with positive surgical margins (Supplementary Table 6). However, there were no significant differences in VPRBP expression between patients with no regional lymph node metastasis (N0) and patients with metastasis (N+). Significant correlations were also not seen between VPRBP expression and pre-operative PSA levels (Supplementary Table 6). To determine whether VPRBP expression associated with poor prognosis disease, we evaluated its expression versus post-operative biochemical (PSA) recurrence-free survival. We observed a statistically significant reduction in PSA recurrence-free survival in patients with any VPRBP expression compared to those who were negative (Fig. S6D). We went on to assess the co-expression of VPRBP with AR or OGT in the TMA. For both, we observed a statistically significant positive correlation in staining which suggests that the expression and activity of these proteins may be relevant in subsets of patients (Fig. 5A and Fig. 5B). Furthermore VPRBP positive/AR high groups exhibited a statistically significant reduction in PSA recurrence-free survival when compared to VPRBP negative/AR low groups (p=0.0019); and so did VPRBP positive/OGT high groups compared to VPRBP negative/OGT negative (p=0.0013) (Fig. 5C and Fig. 5D). We next sought to determine whether VPRBP transcript expression in clinical samples is associated with activity gene signatures reflecting AR activity. First we compared the VPRBP mRNA expression in the TCGA datasets and found a positive correlation with AR mRNA expression (Fig. 5E). There have been multiple studies reporting AR activity gene signatures. In order to identify the subset of AR activity gene signatures which are highly expressed in VPRBP high tumors, we combined AR activity gene signatures from Dorothea AR regulon, KEGG prostate cancer pathway and an additional 8 gene sets from the Molecular Signatures Database (MSigDB) and clustered them on VPRBP high and low expression quartiles as a heatmap (Combined AR gene list in supplementary file 4) (Fig. S7). These gene sets included Nelson androgen stimulated down (17 genes), Nelson androgen stimulated up (83 genes)^36^, PID AR non-genomic (31 genes), PID AR pathway (61 genes), PID AR TF (53 genes), TAKAYAMA bound by AR (10 genes)^37^, Wang response up (28 genes) ^38^ and Wiki pathway androgen pathway signaling (91 genes). Combined, these studies have attributed the expression of 467 genes to AR activity. Since it is not clear which may be most relevant in a given patient group we assessed all of them relative to VPRBP quartile expression in the TCGA Prostate Cancer dataset (Fig.S7). 139 of these 467 genes positively correlated with VPRBP expression and 92 negatively correlated with VPRBP expression (Supplementary file 5). The association between VPRBP expression quartiles and the 139 genes is illustrated in a box plot and is statistically significant between the quartiles Fig. 6F. Importantly the 139 genes most significantly associated with AR responsive pathways but did not include some canonical AR targets such KLK2 and KLK3. In order to further refine this association, we generated co-expression tables for the TCGA dataset ranking all genes that were co-expressed with VPRBP according to a Spearman rank correlation coefficient. We also did this for two other datasets, SU2C/PCF Dream Team, Cell 2015 and MSKCC, Cancer Cell, 2010 from cBioPortal. 48 genes amongst the 139 genes identified were significantly co-expressed with VPRBP (Spearman rank coefficient threshold >0.4) in all the three studies (Supplementary file 5). This represents putative VPRBP-dependent AR regulome which will form the basis for future mechanistic studies on the interplay between VPRBP and AR activity. Next we looked at the p53 activity signatures and found an inverse correlation of VPRBP expression with HALLMARK_P53_pathway ^39^ (Fig. 6G). We also evaluated VPRBP expression versus a second gene signature reflecting p53 activity derived from perturbation experiments (progeny) ^40^ in the Taylor datasets. This also showed a statistically significant inverse correlation (Fig. 6H). Together these studies underscore the significance of VPRBP in promoting PCa growth and progression.

**Figure 5.**
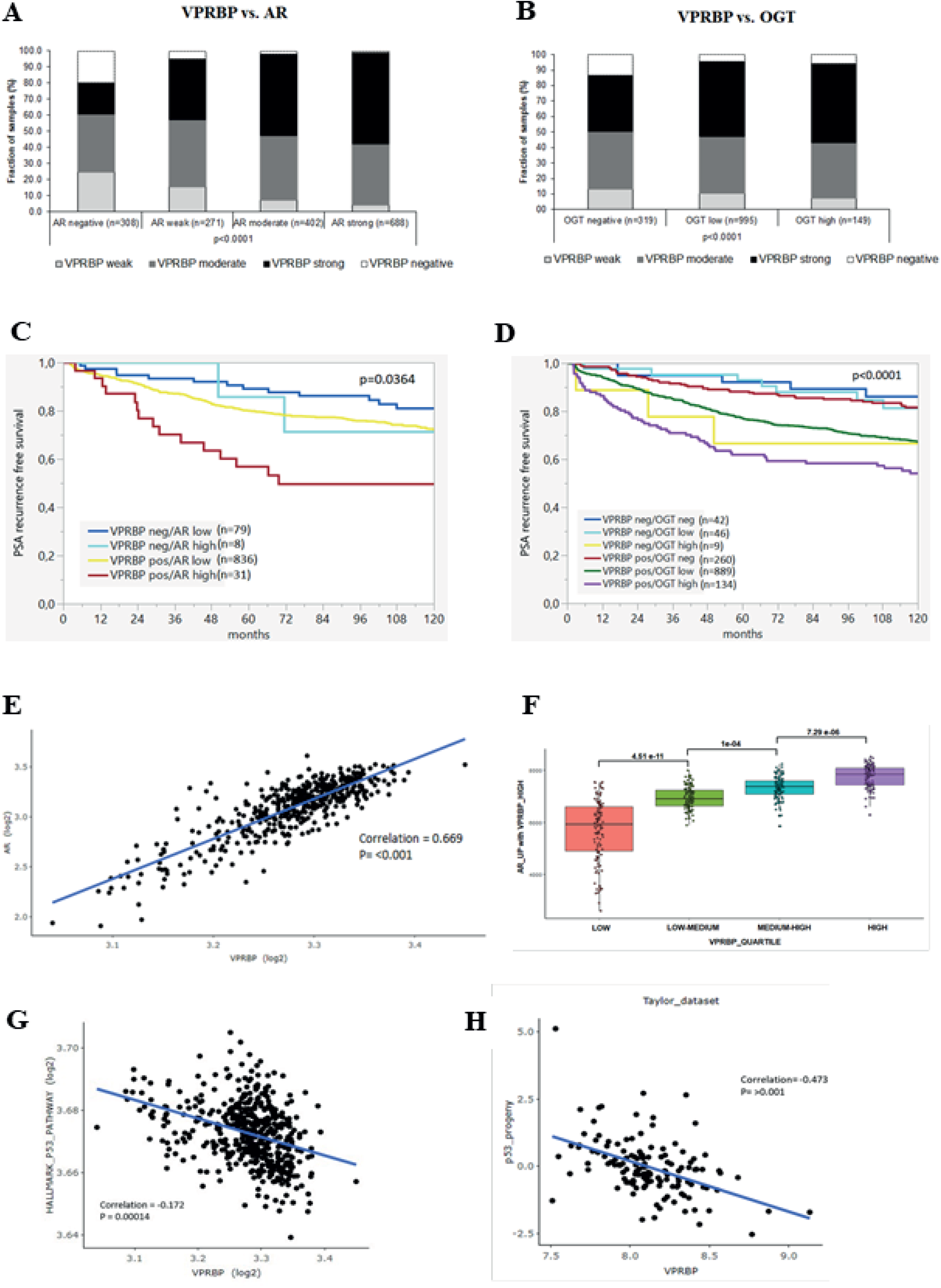
VPRBP protein expression correlates with AR amplification, OGT overexpression and poor prognosis. Bar graphs showing positive correlation of VPRBP expression with AR (A) and OGT expression (B) by IHC in TMA sections. (C) PSA recurrence free survival curves in patients expressing low or high levels of AR where VPRBP expression was present or absent and (D) survival curves in patients expressing no, low or high levels of OGT where VPRBP expression was present or absent. (E) Scatter plot comparing AR mRNA expression to VRPBP mRNA expression in TCGA PanCancer Atlas prostate dataset. (F) Box plot showing association between VPRBP expression quartiles and the 139 up-regulated AR genes from heat map. (G) Scatter plot comparing GSEA Hallmark “P53 Pathway” to VRPBP mRNA expression in TCGA PanCancer Atlas prostate dataset. (H) Scatter plots showing VPRBP expression in Taylor data sets versus p53 progeny scores.

**Figure 6.**
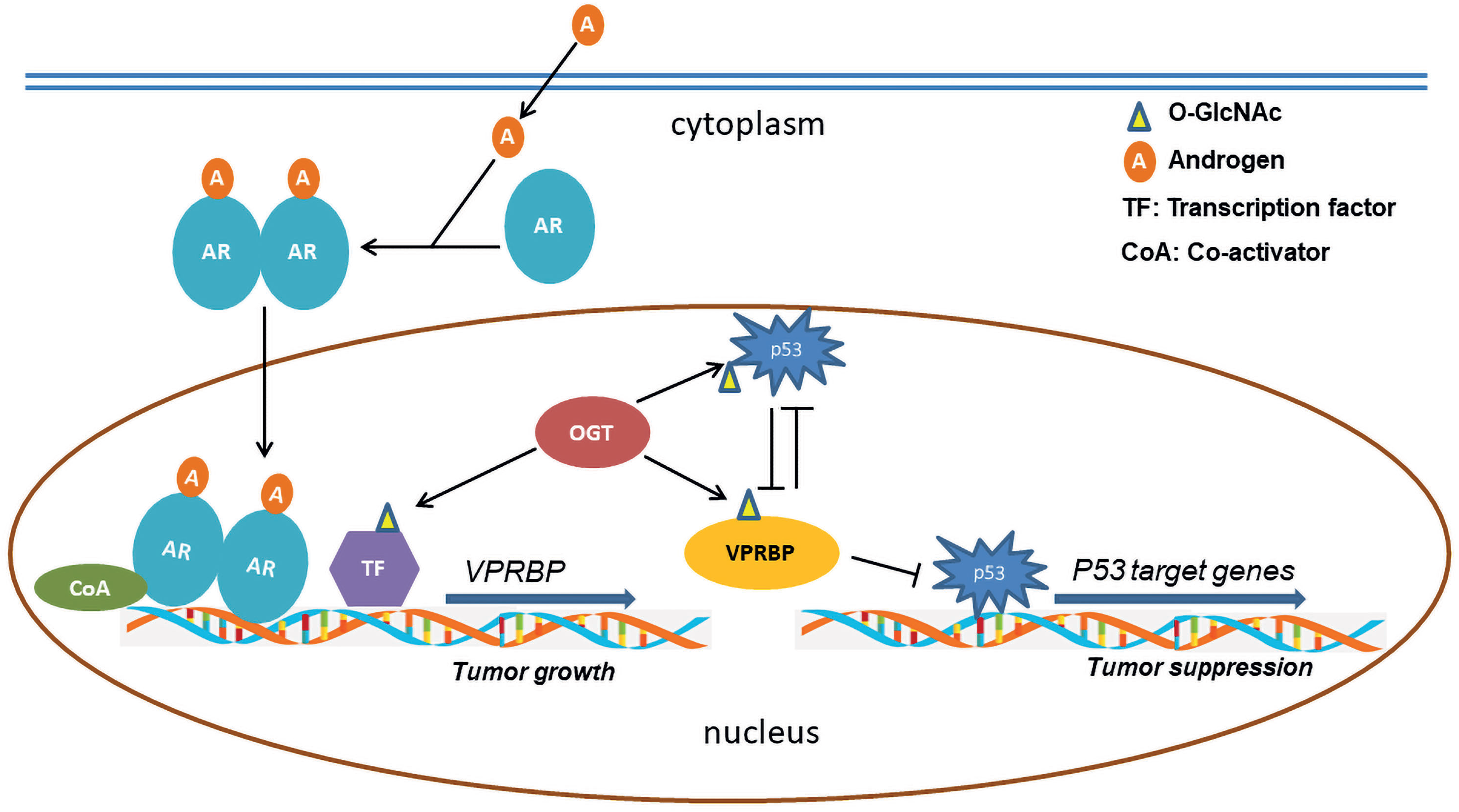
Model depicting VPRBP regulation in prostate cancer. AR activation by androgens (A) stimulate *VPRBP* gene expression in the presence of hitherto unknown O-GlcNAcylated TF/s. VPRBP protein is further stabilized by OGT whereas VPRBP diminishes p53 stability and vice versa. VPRBP also suppresses p53 recruitment to the chromatin thereby regulating p53 activation and inducing cell proliferation and tumor growth.

## Discussion

Our previous studies attributed the impact of O-GlcNAc/OGT on PCa cells to effects on c-Myc stability ^2^, and based on ChIP-seq data, to the over-representation of c-Myc at O-GlcNAc binding sites in the genome ^10^. Though the majority of O-GlcNAc peaks in the genome are promoter proximal and associated with histone marks indicative of active transcription ^10^, the majority of AR sites are distal intergenic or intronic. However, we observed that a small number of AR binding sites overlapped with O-GlcNAc sites on chromatin and hypothesised that these might help us to identify factors that were co-regulated by the AR and OGT. The potential biological significance of these sites is underscored by the fact that in our case they have led us to uncover VPRBP as a novel AR and OGT target.

VPRBP was initially identified as a protein targeted by HIV-1 viral protein R (Vpr) to initiate host cell response leading to cell cycle arrest at G2/M by hijacking the CUL4A E3 ubiquitin ligase machinery ^31 41^. This implies that VPRBP can in some settings support cell cycle progression and indeed we know this to be the case from previous studies ^42^. We show that VPRBP is transcribed in response to androgen treatment (Fig. 1D) and that its protein stability is dependent on OGT (Fig. 2B and Fig. 2E). We went on to show that knockdown of VPRBP by siRNA led to significant decrease in LNCaP proliferation accompanied by stabilization of tumor suppressor p53 (Fig. 3C). Guo *et al* ^12^ demonstrated similar stabilization of p53 in T cells following VPRBP deletion suggesting its requirement in Mdm2-mediated p53 poly-ubiquitination ^12^. They further show that for T cell proliferation to occur, VPRBP promotes cell cycle entry by restraining p53 activation whilst a VPRBP-dependent, p53-independent programme possibly involving c-Myc dictates cell growth in naive T cells after T-cell receptor (TCR) activation. ^12^. One can draw some similarities between T cells progressing from quiescence to proliferation and prostate cancer cells during androgen stimulation. During TCR activation, cells undergo drastic metabolic changes with increased glucose and glutamine uptake, and a concomitant increase in O-GlcNAcylation ^43^. Comparably, androgen stimulation of PCa cells increased glucose uptake and anabolic synthesis of glutamine ^11^. Androgen stimulated cells also displayed higher HBP pathway enzymes and protein O-GlcNAcylation levels (Fig. 1E). Interestingly, VPRBP is upregulated upon TCR activation in T cells as well as with AR activation in PCa cells. Collectively our data suggest that the similar dependencies on both VPRBP and OGT for cell proliferation exist for both PCa cells and T cells.

Other than stabilization, the regulation of p53 transcriptional activity by VPRBP has been previously described by Kim *et al* ^44^ to occur at the chromatin level. In this study they showed that VPRBP is recruited to target promoters by p53 to attenuate p53 dependent transcription by selectively binding to the unacetylated histone H3 tails in the absence of any stress stimuli rendering it inaccessible to Histone acetyl transferases. They also showed that VPRBP knockdown led to activation of p53 target genes, so did its phosphorylation at ser-895 by DNA-activated protein kinase (DNA-PK). A follow up study by the same group further identified a novel intrinsic kinase activity of this protein towards histone H2A on threonine 120 which favours its localization to tumor suppressor genes and chromatin silencing ^15^. In our study we have shown that VPRBP knockdown significantly enhances the recruitment of p53 to chromatin as assessed by a significant increase in genome-wide p53 binding sites. Collectively, these suggest that VPRBP is a multi-stage inhibitor of p53 activation, impacting on both chromatin binding and p53 stability and expression. This impact may however be most profound in cells expressing wild-type p53 since a mutant p53 cell-line, VCaP, did not show significant reductions in p53 levels with VPRBP knockdown (Fig. 3E). We also tested the feedback effects of p53 activation on VPRBP expression and have found that stabilising p53 pharmacologically with nutlin-3a diminishes VPRBP stability and knocking out p53 enhances it (Fig. S4F). We believe this is predominantly a post-translational/protein turnover effect since there are no significant changes in transcript levels (Fig. S4G) and no evidence of p53 binding to the VPRBP promoter (Figure S4E). All the above studies suggest a reciprocal relation between p53 and VPRBP in PCa cells.

We have also shown previously that inhibiting guanine nucleotide biosynthesis disrupts nucleolar function leading to p53 stabilization and c-Myc downregulation ^16^. In that study we reported that inhibiting IMPDH2 with a drug, mycophenolic acid, led to p53 stabilisation by depleting cellular GTP levels, promoting degradation of nucleolar proteins such as GNL3 and thereby inducing nucleolar stress ^16^. Interestingly mycophenolic acid was developed and used initially to restrict T and B cell proliferation for the purposes of enhancing graft take in patients undergoing renal transplant surgery ^45^. This further reinforces the idea that there are significant commonalities in the biological processes that support immune activation mediated T-cell proliferation and PCa cell proliferation. Since VPRBP was previously shown to sustain 40S ribosome subunit biogenesis by supporting nucleolar integrity ^30^, we were led to test how targeting VPRBP might affect this compartment. Imaging of nucleolar marker, fibrillarin revealed marked changes in nucleolar staining indicative of nucleolar stress (Fig. 4G). A recent study by Han *et al* revealed a critical role for VPRBP in rRNA processing and ribosome biogenesis by regulating a previously unknown substrate, the ribosome assembly factor PWP1 ^13^. Interestingly, VPRBP loss leads to accumulation of free ribosomal protein L11 (RPL11), resulting in L11-MDM2 association and p53 activation. Together, we conclude that VPRBP restricts p53 activation in part by maintaining nucleolar integrity.

By examining the VPRBP protein expression in tissue from a highly annotated PCa patient cohort, we established that expression increases significantly with stage and grade and furthermore correlates positively with high expression of the AR and OGT. Future studies will also need to further dissect the functional impact of VPRBP in a range of other mutational backgrounds include RB-loss, PTEN-loss and p53 point mutation. In conclusion VPRBP represents the first AR and OGT co-regulated protein to promote prostate cancer cell proliferation by limiting p53 activation and as such may be an early determinant of prostate cancer progression (Findings summarized in Fig.6). Based on our studies and previous studies on T cell proliferation and activation we believe that it works hand-in-glove with c-Myc to support proliferation. It would be relevant to include VPRBP in patient stratification for treatment optimization in men with PCa.

## Materials and Methods

### Reagents and consumables

Synthetic androgen, R1881 and dihydrotestosterone were obtained from Sigma-Aldrich. OGT inhibitors, OSMI2 and OSMI3 were kindly provided by Professor Suzanne Walker (Harvard Medical School, Boston, MA, USA). Formaldehyde 16% (F017/3) was purchased from TAAB laboratory. iDeal ChIP-seq Kit for Transcription Factors (C01010170) was obtained from Diagenode. B32B3 (SML1419) was purchased from Sigma-Aldrich. Antibody details are provided in supplementary table 1. Lipofectamine RNAiMAX Transfection Reagent (13778075) was obtained from ThermoFisher scientific. Protein A sepharose beads (ab193256) and protein G sepharose beads (ab193259) were from abcam.

### Cell lines

LNCaP cells were purchased from ATCC and cultured in Roswell Park Memorial Institute media (RPMI) containing 10% fetal bovine serum and 1% pencillin-streptomycin in a humidified incubator at 37°C and 5% CO_2_. VCaP cells were obtained from ATCC and cultured in Dulbecco’s Modified Eagle Media (DMEM) containing 10% fetal bovine serum and 1% pencillin-streptomycin in a humidified incubator at 37°C and 5% CO_2_. TP53 CRISPR knockout LNCaP cells were provided by Dr.Simon McDade (Queen’s University, Belfast, UK; supplementary materials and methods)

### Chromatin immunopreciptation (ChIP) assays

ChIP assays were carried out as per manufacturer’s instructions (Details in supplementary materials and methods).

### Real time qPCR

The cells were lysed in qiazol and RNA isolated using Qiagen miRNeasy mini kit (Cat # 217004). Transcriptor first strand cDNA synthesis kit (Cat # 04897030001, Roche Life science) was used for cDNA preparation. SYBR Green 1 Master (Cat# 4887352001, Roche Life science) was used to compare gene expression changes in VPRBP, OGT, CAMKK2, UAP1, COPS3, p53 and p21 by realtime qPCR in Roche LightCycler® 480 Instrument II. Human large ribosomal protein (RPLPO) was used as the internal control. Primers for realtime PCR were purchased from either Sigma (KiCqStart predesigned) or from Eurofins genomics. The primer details are provided in supplementary table 3 and 4 respectively.

### siRNA transfection

LNCaP cells were seeded on to 6 well plates for siRNA knockdown. Forward transfection was performed the following day using Lipofectamine RNAiMAX Transfection Reagent in OPTI-MEM with 30 pmol of siRNA/well according to the manufacturer’s instructions. After overnight incubation, media was changed to RPMI with 10% FBS and antibiotics. Two individual siRNAs were used against each target of interest. The siRNAs used were OGT si1, OGT si2, DCAF1 si1, DCAF1 si2, and Silencer™ Select Negative Control No.1 siRNA (Details are provided in supplementary table 5). For studies involving androgen treatment, media was changed to androgen deprived charcoal-treated media for 3 days prior to stimulation with 1 nM R1881 for 24h. The cells were lysed after given number of days post transfection for analysis by western blot or real time PCR.

### Western blot analysis

The cells were lysed in RIPA buffer (Sigma) containing protease inhibitor cocktail (Roche) and PhosSTOP™ (Sigma); protein concentration estimated with Bradford reagent (Biorad) and 30 μg of lysate subjected to electrophoresis using precast 4–12% NuPage mini-gels (Life Technologies). The resolved proteins were then transferred to PVDF membrane, blocked with 5% nonfat dry milk and probed with respective primary antibodies overnight at 4°C. After 3 washes, the blots were probed with HRP-conjugated secondary antibodies and immunoreactivity detected by enhanced chemiluminescence in Syngene G box.

### Immunoprecipitation

Briefly, LNCaP cells were lysed in IP lysis buffer (10mM Tris-Cl pH7.5, 140mM NaCl, 1mM EDTA, 1% Triton X-100, 0.1% sodium deoxycholate, 0.1% SDS). Lysates were precleared with protein A sepharose beads for 1h at 4°C in a rotator. Immunoprecipitation (IP) was carried out with protein A sepharose beads using 1mg lysate and 1 μg VPRBP antibody. Lysate was incubated with VPRBP or IgG negative control antibody for 3h at 4°C in a rotator followed by overnight incubation with 40μl of washed protein A sepharose beads. The beads were briefly pelleted and washed thrice in IP wash buffer (10mM Tris-Cl pH7.5, 150mM NaCl, 0.5% Triton X-100). The proteins were eluted by heating in 20 μl laemelli buffer at 95°C for 5min.

### Cell counts

Cell counting of siRNA transfected cells in 12 well plates was performed 5d post transfection. Cells were trypsinized and changes in cell number assessed by cell counting in Countess II Lifetechnologies.

### Immunoflourescence

LNCaP cells were seeded on to glass coverslips in a 12 well plate. After reaching 70-80% confluence, cells were transfected with scrambled or VPRBP siRNA. After 3 days, cells were fixed with 4% paraformaldehyde in BSA for 10 min followed by three washes in PBS for 5 min each. The fixed cells were then lysed in 0.1% TX100 in PBS for 10 min and blocked in blocking buffer (5% goat serum/1%BSA/0.1%TX100 in PBS) for 1h. The cells were then incubated in primary antibodies against VPRBP (1:80) and fibrillarin (1:100) for 2h at RT followed by Alexa flour 594 Goat anti mouse (Invitrogen, Cat # A11020) or Alexa flour 488 Goat Anti R (Invitrogen, Cat # A11070) secondary antibodies. Coverslips were mounted on to glass slides using Vectashield with DAPI.

Patients, immunohistochemistry and analysis of TCGA and other datasets can be found in supplementary materials and methods

## Statistical analysis

Statistical analyses for studies in LNCaP and VCaP cells were done using either student’s t-test or one way ANOVA with Tukey’s posthoc analysis, as mentioned in the figure legends. For IHC, statistical calculations were performed using JMP 12® software (SAS Institute Inc., NC, USA). Contingency tables were calculated with the chi^2^-test. Survival curves were calculated by the Kaplan-Meier method and compared with the Log rank test.

## Acknowledgments

This research was supported by the Norwegian Research Council (230559). IGM is also supported by the Prostate Cancer UK/ Movember Centre of Excellence (CEO13_2–004) and the John Black Charitable Foundation. Authors would like to acknowledge Genomic Core Technology Units, Queens University Belfast for the assistance with ChIP-seq studies.

## Author contributions

NP designed and performed the experiments, prepared samples for sequencing, analyzed the data and wrote the manuscript. AP, GS and Sarah Minner performed IHC and evaluated expression in prostate tumours. NF performed analysis on TCGA, Taylor and other datasets from MSigDB. GG generated p53 knockout LNCaP cells under the supervision of SMcD. SMcD reviewed and edited the manuscript. MF performed the fragment analysis for ChIP-seq samples and conducted the Illumina NextSeq™ 500 sequencing run. Sarah Maguire generated the fastq files, bam files and bed files. IGM supervised the study, reviewed and edited the manuscript.

## Supplementary figures

**Supplementary Figure S1.**
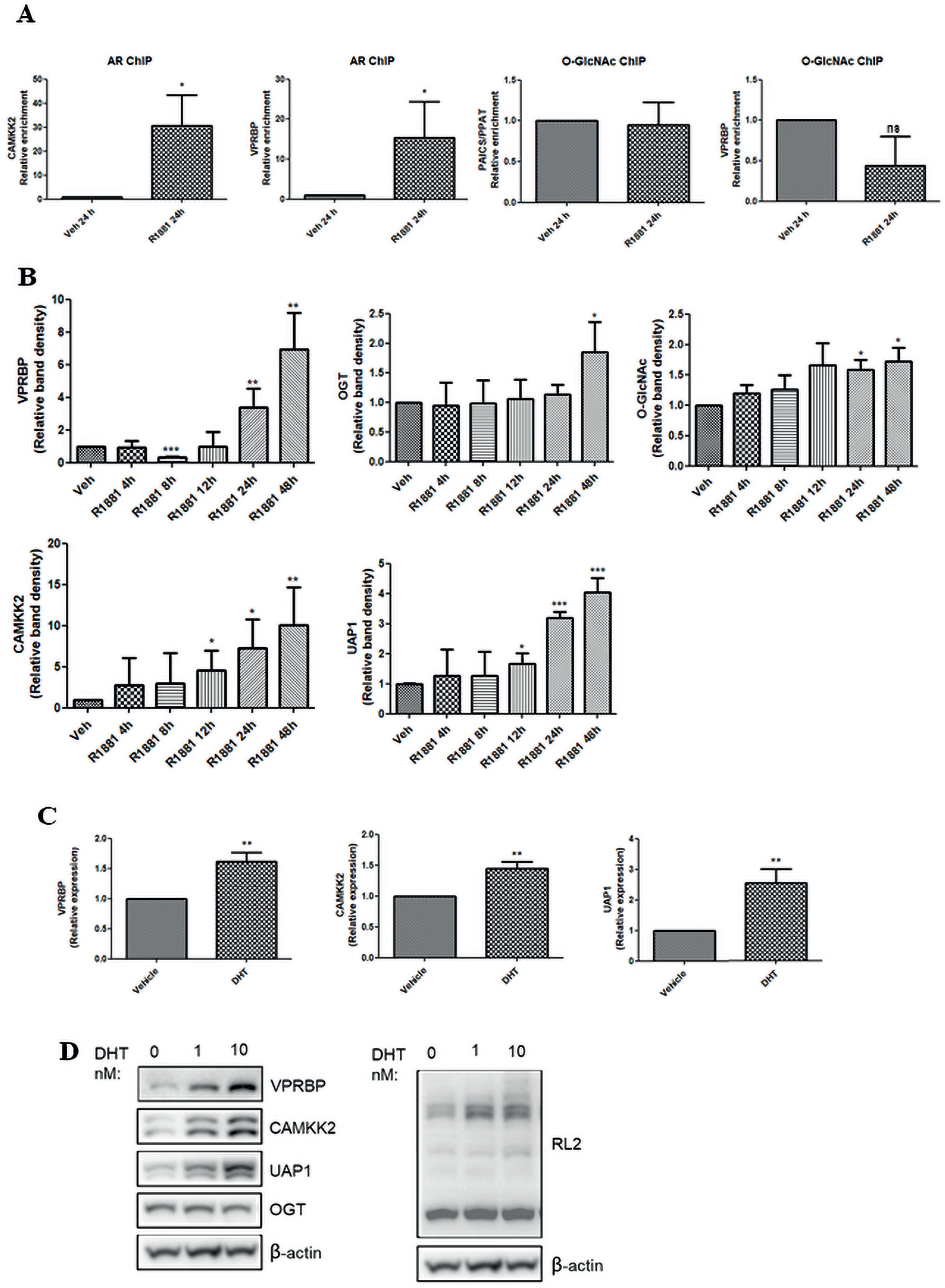
VPRBP is a novel AR target (A) Bar graphs showing relative enrichments in CAMKK2 and VPRBP with AR ChIP-qPCR in vehicle vs R1881 (24h) treated LNCaP cells; and relative enrichments in PPAT/PAICS and VPRBP with O-GlcNAc ChIP. (B) Graphs showing quantitation of western bands of Figure 1E done using image J software and normalized to β-actin. (C) qRT-PCR analysis of VPRBP, CAMKK2 and UAP1 expression in LNCaP cells treated with vehicle (0.01% ethanol) or 1nM DHT for 24h following androgen deprivation for 72h. (D) Western blot analysis of LNCaP cells treated with vehicle or DHT (1nM and 10nM) for 24 h. Results are expressed as means ± SD. * =p < 0.05, ** =p < 0.01; *** = p < 0.001 by Student’s t test.

**Supplementary Figure S2.**
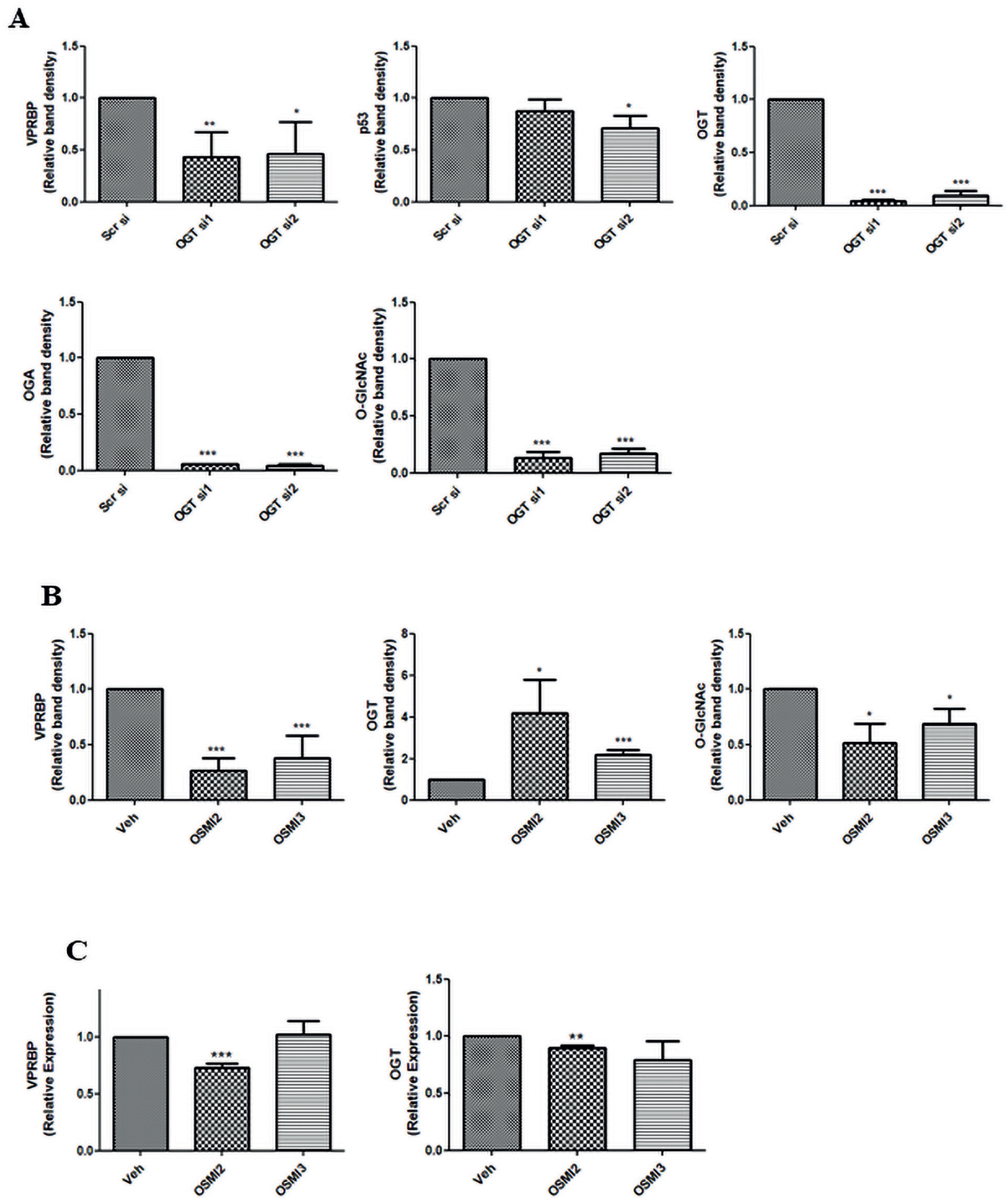
OGT is required for VPRBP stability. (A) Graphs showing quantitation of western bands (Fig. 2B) from OGT siRNA transfected cells vs scrambled control using image J software and normalized to β-actin. (B) Graphs showing quantitation of western bands (Fig. 2E) from OSMI2/OSMI3 treatments and normalized to β-actin. (C) qRT-PCR analysis VPRBP and OGT expression in LNCaP cells treated with vehicle or 40 μM OSMI2 or 10 μM OSMI3 for 24h with RPLPO as internal control. Results are expressed as means ± SD. * =p < 0.05, ** =p < 0.01; *** = p < 0.001 by Student’s t test.

**Supplementary Figure S3.**
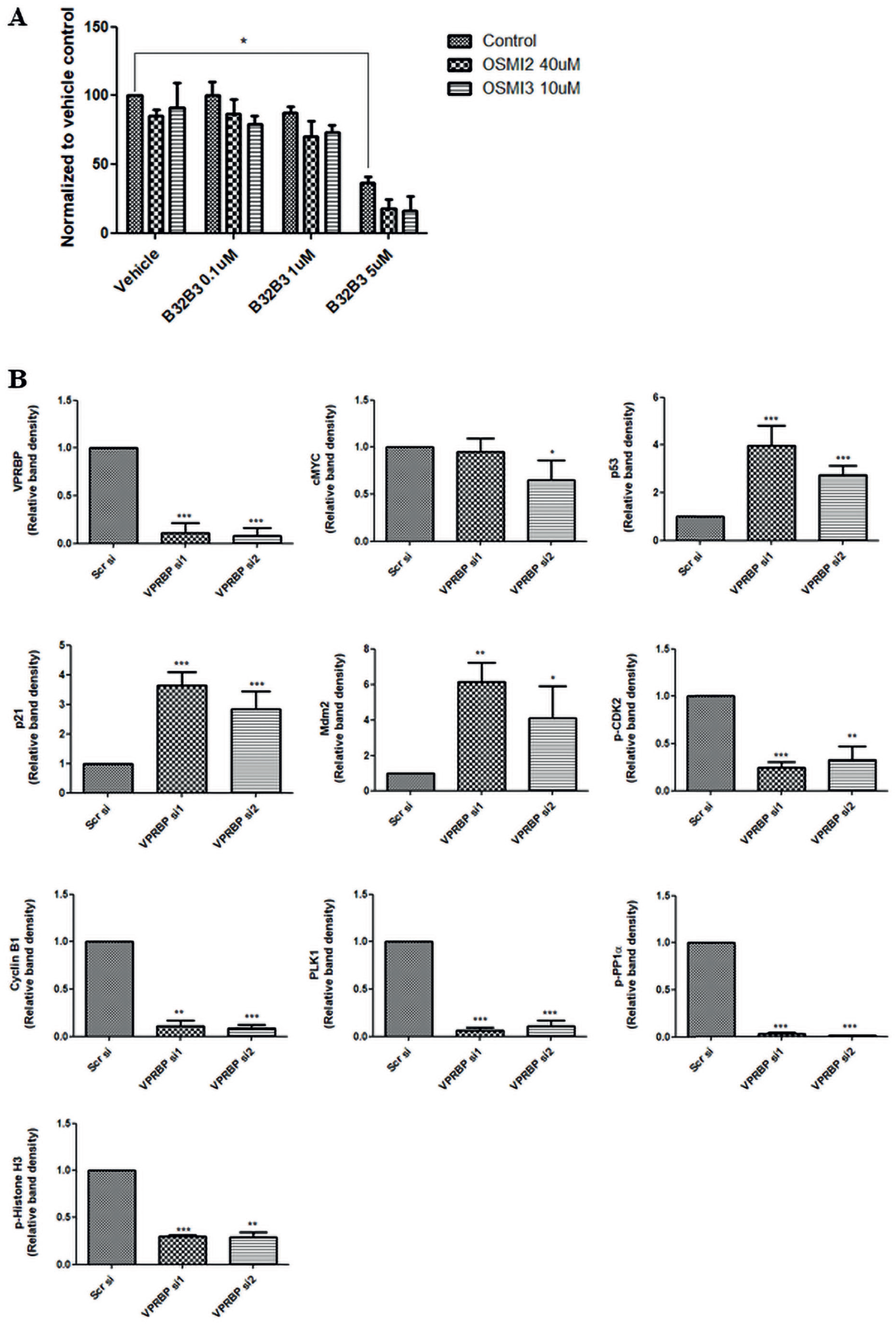
(A) Effect of VPRBP kinase activity inhibitor, B32B3 on LNCaP cell proliferation was assessed by cell counting 4d post treatment in the presence and absence of OSMI2 and OSMI3. (B) Graphs showing quantitation of western bands (Fig. 3C) from VPRBP siRNA transfected cells *vs* scrambled control using image J software and normalized to β-actin. Results are expressed as means ± SD. * =p < 0.05, ** =p < 0.01; *** = p < 0.001 by Student’s t test.

**Supplementary Figure S4.**
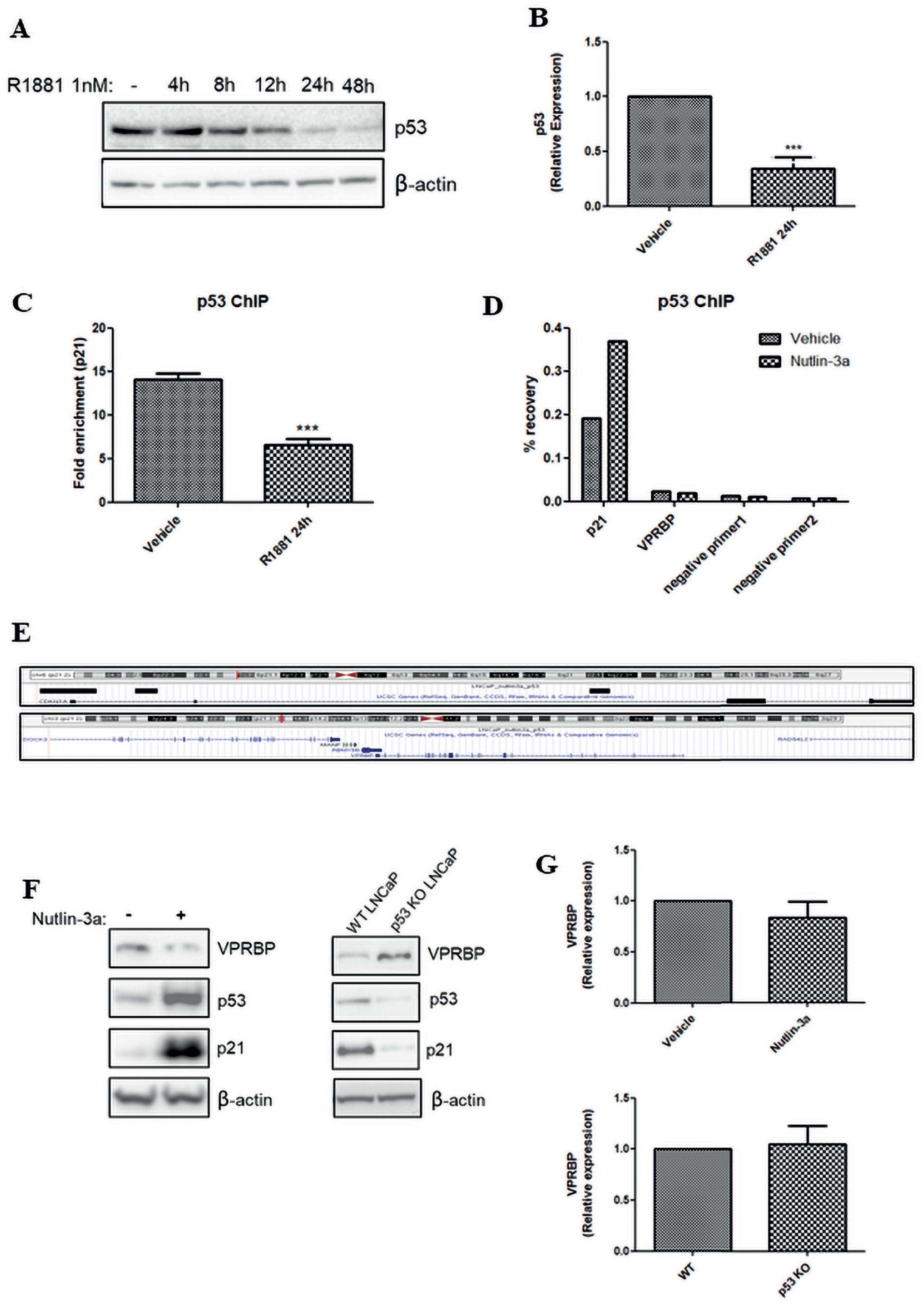
VPRBP and p53 exhibit reciprocal relation. (A) Western image showing time dependent reduction in p53 expression following exposure to 1nM R1881 for different time points. (B) qRT-PCR analysis of p53 expression in LNCaP cells treated with vehicle or 1nM R1881 for 24h with RPLPO as internal control. (C) Bar graphs showing relative enrichments in p21 with p53 ChIP-qPCR in vehicle and R1881 24h treated LNCaP cells. (D) Bar graphs showing percentage recovery of p21, VPRBP and negative primers with p53 ChIP-qPCR in vehicle (DMSO) and nutlin-3a 24h treated LNCaP. (E) ChIP-seq enrichment profile of nutlin-3a p53 ChIP-seq at CDKN1A (p21) promoter region (top) and absence of enrichment at VPRBP promoter region (bottom), using UCSC genome browser. (F) Protein expression of VPRBP, p53 and p21 by immunoblot analysis following vehicle and nutlin-3a treatment for 24h; and wild type versus p53 knockout LNCaP cells, and (G) mRNA expression by qRT-PCR. Results are expressed as means ± SD. *** = p < 0.001 by Student’s t test.

**Supplementary Figure S5.**
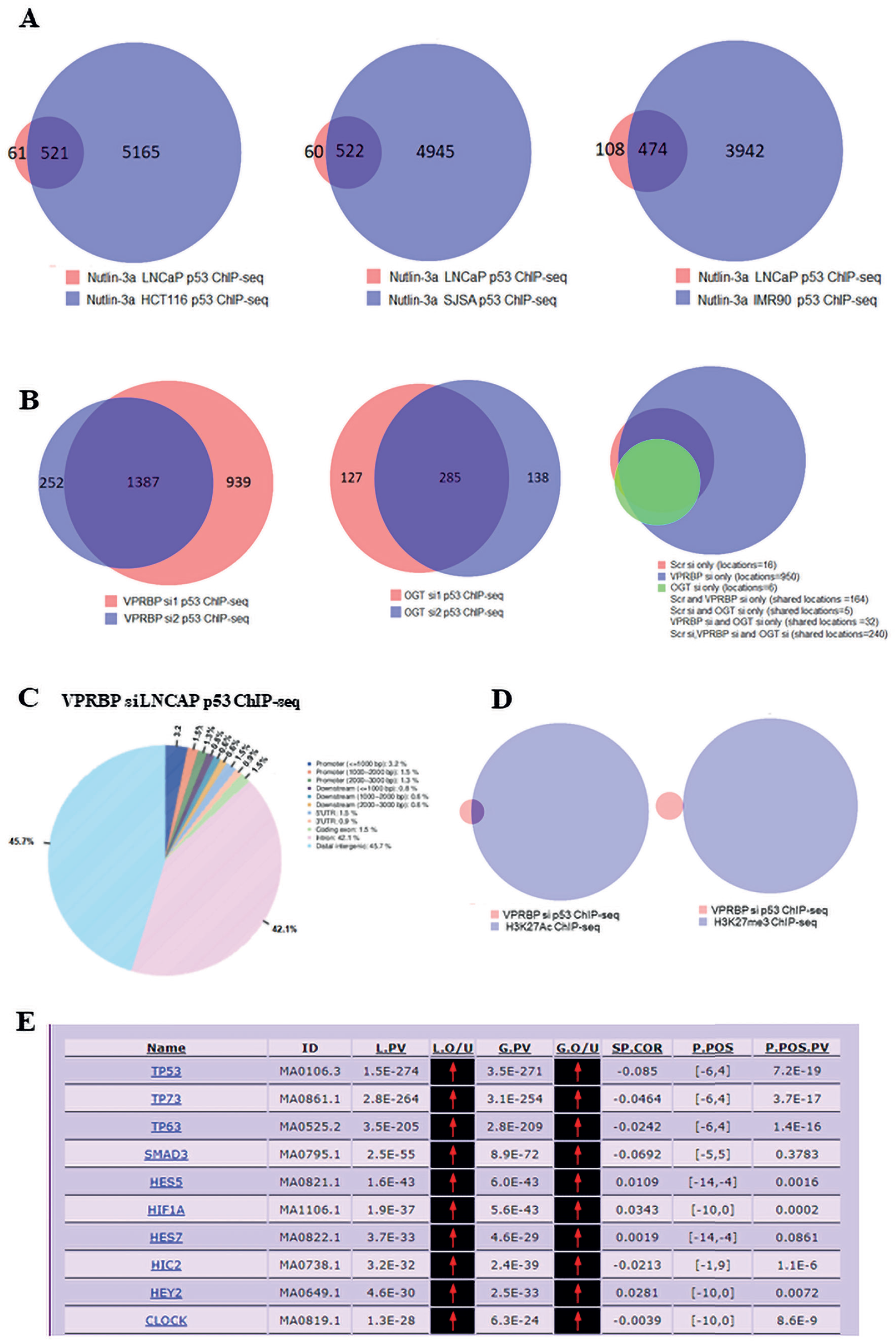
(A) Venn diagram showing the overlap of our nutlin-3a p53 ChIP-seq in LNCaP with previously reported nutlin-3a p53 ChIP-seq in HCT116 (GSE86164), Bone/SJSA (GSE86164) and IMR90 fetal lung fibroblasts (GSE58740). (B) Venn diagram showing the overlap of VPRBP si1 and VPRBP si2 p53 ChIP-seq peaks; overlap of OGT si1 and OGT si2 p53 ChIP seq peaks; and overlap of scr, VPRBPsi and OGTsi ChIP-seq peaks. (C) CEAS analysis of VPRBP si p53 ChIP-seq in LNCaP. (E) Venn diagram showing overlap of VPRBP si p53 ChIP-seq with H3K27Ac ChIP-seq and H3K27me3 ChIP-seq in ethanol treated LNCaP-MYC cells (GSE73994) (F).Table showing the top over-represented motifs obtained from motif enrichment analysis of VPRBP si p53 ChIP-seq using P-scan ChIP.

**Supplementary Figure S6.**
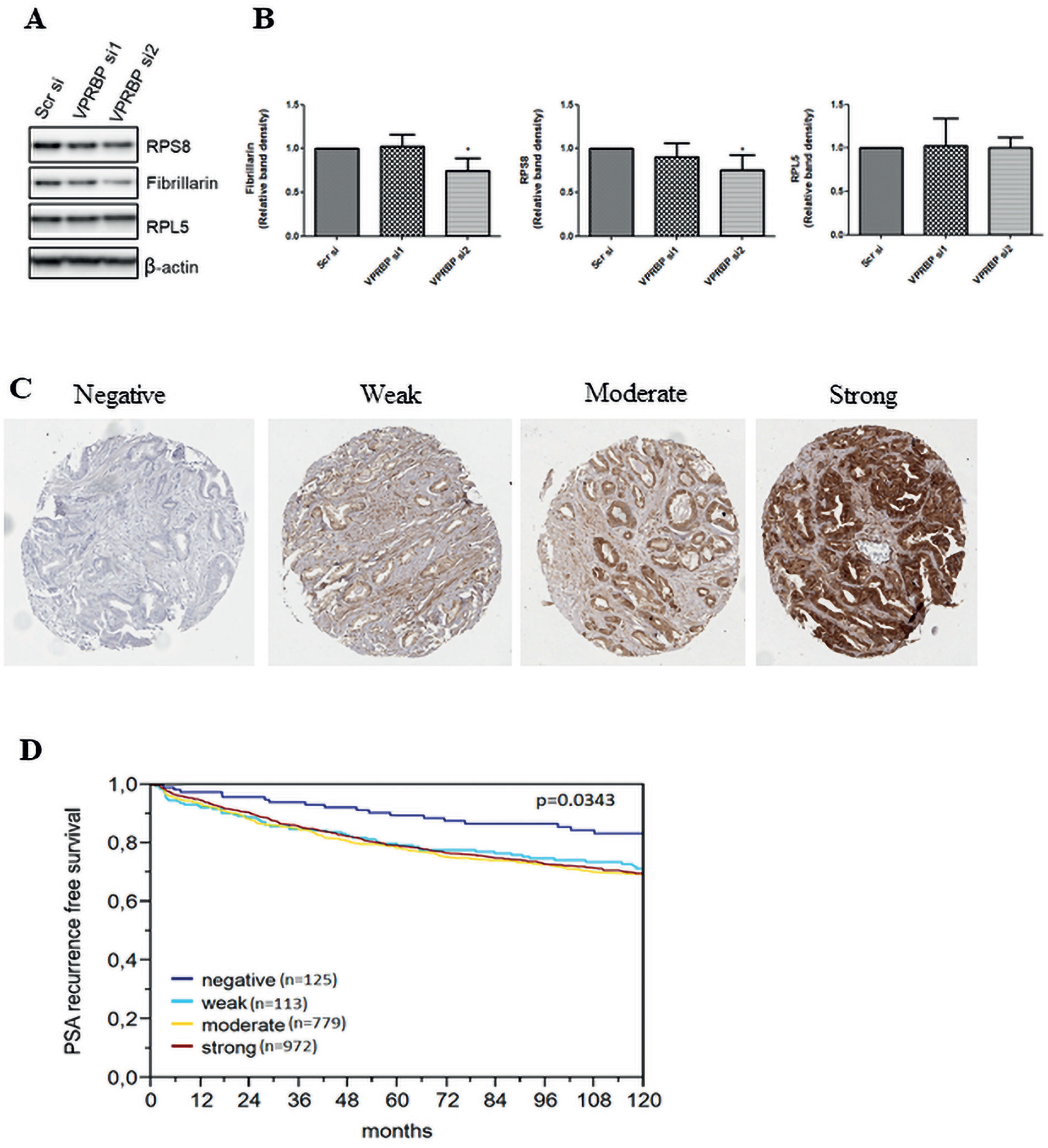
(A) Effect of VPRBP knockdown on fibrillarin, RPS8 and RPL5 was assessed by immunoblot analysis and (B) graphs showing quantitation of the western bands. (C) IHC representative images showing negative, weak, moderate and strong staining of VPRBP. (D) Survival curve showing inverse relation between VPRBP expression and PSA recurrence free survival. Results of western quantitation are expressed as means ± SD. * =p< 0.05 by Student’s t test.

**Supplementary Figure S7.** Heatmaps comparing the expression of a total of 467 AR activity gene signatures combined from multiple studies, in high and low VPRBP quartiles, created using the R package.

