## Supplementary materials methods and tables for "VPRBP functions downstream of the androgen receptor and OGT to restrict p53 activation in prostate cancer"

**Supplementary Materials and Methods**

**Generation of CRISPR-Cas9 Knockout LNCaPs**

Chemically competent DH5α was prepared by standard calcium chloride method and transformed with pD1301-AD vector containing the gene encoding the Cas9 nuclease. Colonies with incorporated plasmid were selected using kanamycin resistance and glycerol stocks were subsequently prepared from colonies grown in LB media. The remaining culture was used to purify plasmid DNA by mini prep using Zyppy™ Plasmid Miniprep Kit by (Zymo Research) according to manufacturer's instructions. Guide RNA sequences for TP53 were sourced from Horizon Discovery- plasmid map and can be found in the attached figure and table. For transfecting CRISPR plasmid DNA, LNCaP cells were seeded in a Nunc™ 6 well tissue culture plate at a density of $3.0 \times 10^5$/mL. After 24 hours, 1μg of plasmid DNA was combined with 3μL of Fugene HD transfection reagent (Promega) and optiMEM (Gibco) to a final volume of 100μL and added to cells replenished with 1mL of fresh culture medium. After 24 hours, successful transfection of plasmid DNA was investigated using an EVOS FLoid Cell Imaging Station (Thermo Fisher Scientific) and pictures recorded. Provided there was successful plasmid uptake, the cells were replaced with media for a further 24 hours. Cell pools were expanded from 6 well culture plates into T-25cm$^2$ culture flasks and allowed to adhere before 5μM of Nutlin-3a was supplemented into the media as a selection agent, thereby positively selecting for those cells with loss of function of TP53. The cell pools remained in selection until total death of cells in an untransfected control flask was evident. Following selection of the LNCaP cells with Nutlin-3a, cells were left as a knockout pool due to the inability this cell line to efficiently form single cell colonies.

A

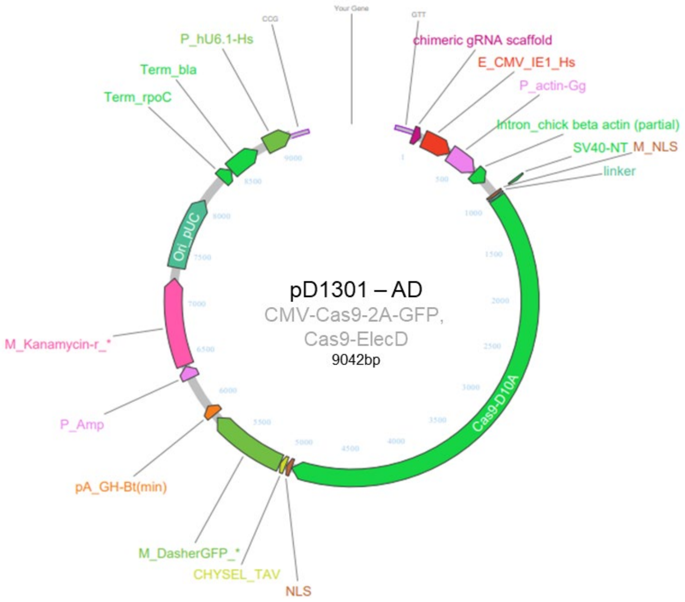

B

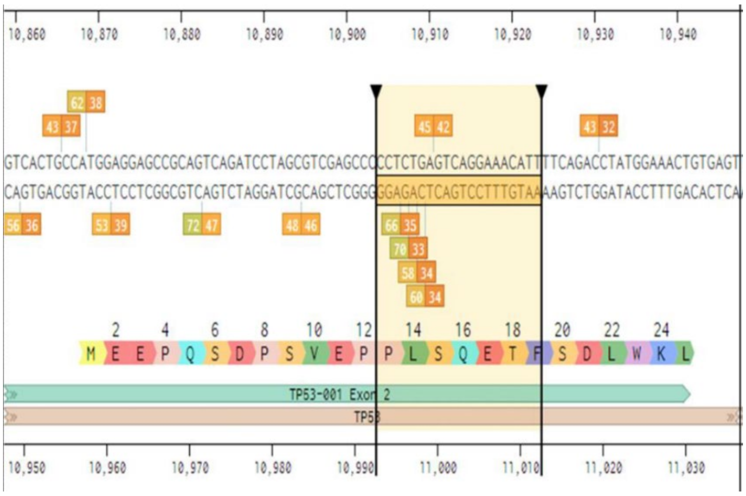

C

| Gene | gRNA sequence (bp) |
|---|---|
| *TP53* sgB | AATGTTTCCTGACTCAGAGG (20bp) |
| *TP53* sgC | GATCCACTCACAGTTTCCAT (20bp) |

**Details of CRISPR Cas9 materials utilised to generate novel isogenic models. (A)** Plasmid map of pD1301 – AD that was used for p53 CRISPR-Cas9 knockouts. **(B)** Diagram depicting the genomic target of *TP53* sgB sequence as exon 2. **(C)** Guide RNA sequences which were utilised in deriving p53 knockout cell lines.

**ChIP-qPCR**

LNCaP cells were cultured in androgen deprivation media for 3 days prior to treatment with R1881 1nM, for 24h. Cells were fixed with 1% formaldehyde and chromatin prepared by sonication in Bioruptor® Plus sonication device (Diagenode). Fragmentation efficiency was analysed using fragment analyser (Agilent Technologies) and chromatin immunoprecipitation (ChIP) carried out using diagenode iDeal ChIP-seq kit for transcription factors (C01010055). Commercially available antibodies targeted against AR, O-GlcNAc modification (RL2) and p53 were used for the IPs. Percentage recovery of immunoprecipitated DNA relative to input was calculated from real time quantitative PCR (ChIP-qPCR) Ct values. ChIP-qPCR primers were obtained from Eurofins genomics and sequences listed in supplementary table 2.

**p53 ChIP-seq**

For the p53 ChIP –seq studies, LNCaP cells were grown in 10 cm dishes and treated with DMSO or 2.5 μM nutlin-3a for 24h. Another set was transfected with scrambled, VPRBP or OGT siRNAs. Cells were processed as described in the previous section. ChIP-qPCR of p21 was conducted to measure the enrichment of immunoprecipitated DNA. Libraries were prepared using diagenode Microplex library preparation kit V2, pooled and sequenced using Illumina NextSeq™ 500 high output, yielding ~50 million reads per sample (NextSeq run metrics table and Multi QC report provided in supplementary file 1). Fastq files were generated with Illumina pipeline software (bcl2fastq version 2.19 using the default thresholds). Reads were mapped to the GRCh37/hg19 reference genome using bowtie2 (version 2.2.6) and subsequently filtered to remove PCR duplicates. Prior to peak calling, Encode curated blacklisted genomic regions were removed from bam files. Fastq files were aligned to hg19 using bowtie2 [1]. The MACS algorithm (version 2.1.2) [2] was used to analyse the resulting alignments and identify transcription factor binding regions. The p53 ChIP-seq data has been deposited in the NCBI GEO data repository (GSE169129).

### ChIP-seq data analysis

Galaxy cistrome [3] and galaxy Europe [4] analysis platforms were used to compare AR, O-GlcNAc and p53 ChIP-seq binding sites. For AR ChIP-seq data (GSE28126), binding sites were converted from hg18 to hg19 using the liftover tool in Galaxy cistrome. Galaxy intersection tool was used to intersect intervals of two datasets to return overlapping pieces of intervals for at least 1bp. Galaxy subtract tool was used to subtract intervals of two datasets. CEAS (Cis-regulatory element annotation system) tool in galaxy cistrome was used to annotate ChIP binding sites and BETA-minus for target prediction. Pscan-ChIP was used for motif enrichment analysis [5]. Venn diagrams were generated either in galaxy cistrome or by Venny 2.1.0.

### Patients

Radical prostatectomy specimens were available from 3261 patients, consecutively treated at the Department of Urology, University Medical Center Hamburg-Eppendorf between 1992 and 2005 (Supplementary Table 6). Follow-up data were available for 2385 patients, ranging from 1 to 144 months (mean, 34 months). None of the patients received neo-adjuvant or adjuvant therapy. Additional (salvage) therapy was only initiated in case of a biochemical relapse (BCR). In all patients, prostate specific antigen (PSA) values were measured quarterly in the first year, followed by biannual measurements in the second and annual measurements after the third year following surgery. Recurrence was defined as a postoperative PSA of 0.1 ng/ml and rising. The first PSA value above or equal to 0.1 ng/ml was used to define the time of recurrence. Patients without evidence of tumor recurrence were censored at last follow-up. All prostatectomy specimens were analyzed according to a standard procedure. All prostates were completely paraffin-embedded, including whole-mount sections as previously described [6]. One 0.6 mm tissue core was punched out from each case, and transferred in a tissue microarray (TMA) format as previously described [7]. The 3261 cores were distributed among 7 TMA blocks each containing 129-522 tumor samples. Each TMA block also contained various control tissues including normal prostate tissue and other normal tissues.

The tissues and clinical data were utilized according to the Hamburger Krankenhaus Gesetz (§12 HmbKHG) and approved by our local Ethical Committee.

**Immunohistochemistry**

Freshly cut TMA sections were stained on one day in a single experiment. High-temperature pretreatment of slides was done in an autoclave in citrate buffer, pH 7.8 for 5 minutes. VPRBP immunostaining was performed using a monoclonal antibody (clone: EPR16012, Abcam; dilution: 1:450). The Envision system (DAKO) was used to visualize the immunostaining. Only cytoplasmatic staining was evaluated. The staining intensity (0, 1+, 2+, 3+) and the fraction of positive tumor cells were recorded for each tissue spot. A final score was built from these two parameters according to the following scores: Negative scores had staining intensity of 0, weak scores had staining intensity of 1+ in ≤70% of tumor cells or staining intensity of 2+ in ≤30% of tumor cells; moderate scores had staining intensity of 1+ in >70% of tumor cells, staining intensity of 2+ in >30% and ≤70% of tumor cells or staining intensity of 3+ in ≤30% of tumor cells and strong scores had staining intensity of 2+ in >70% of tumor cells or staining intensity of 3+ in >30% of tumor cells.

**Analysis of TCGA and other datasets**

TCGA PanCancer Atlas 2014 and Taylor datasets were downloaded from cbioportal. For AR activity gene signatures, AR target genes were pulled from the Dorothea AR regulon (138 genes) and KEGG prostate cancer pathway (89 genes). An additional 8 gene sets were downloaded from the Molecular Signatures Database (MSigDB). Scatterplots showing normalized expression were plotted using R studio. ssGSEA was carried out using Genepattern. Single sample scores were calculated for h.all.v7.2 [Hallmarks] gene set in TCGA samples. Progeny scores were calculated using the progeny tool [8]. Pearson correlation coefficient analysis was used to assess correlation. Heatmaps were created using the R package Pheatmap.

**Supplementary Table 1**

| Antibody | Catalog number | Manufacturer | Application |
|---|---|---|---|
| **Androgen Receptor** | ab74272 | Abcam | ChIP-qPCR |
| **RL2** | ab2739 | Abcam | ChIP-qPCR, western |
| **p-53** | sc-126 | Santa Cruz Biotechnology | ChIP-seq, western |
| **VPRBP** | ab202587 | Abcam | Western, IF |
| **OGT** | ab177941 | Abcam | Western |
| **c-Myc** | ab32072 | Abcam | Western |
| **β-actin** | ab6276 | Abcam | Western |
| **Histone H2A phospho T120** | ab177391 | Abcam | Western |
| **Histone H3 (tri methyl K27** | ab6002 | Abcam | Western |
| **Fibrillarin** | ab4566 | Abcam | Western, IF |
| **CSN2** | ab155774 | Abcam | Western |
| **JAB1/CSN5** | ab12323 | Abcam | Western |
| **RPS8** | ab226361 | Abcam | Western |
| **RPL5** | ab137617 | Abcam | Western |
| **Cyclin B1** | 4138 | Cell Signaling Technology | Western |
| **PLK1** | 4513 | Cell Signaling Technology | Western |
| **phospho-PP1α** | 2581 | Cell Signaling Technology | Western |
| **phospho-Histone H3 (Ser10)** | 9701 | Cell Signaling Technology | Western |
| **phospho-CDK2** | 2561 | Cell Signaling Technology | Western |
| **CDK2** | 2546 | Cell Signaling Technology | Western |
| **p-FoxM1** | 14655 | Cell Signaling Technology | Western |
| **p-21** | 2947 | Cell Signaling Technology | Western |
| **goat anti-rabbit** | 7074 | Cell Signaling Technology | Western |
| **goat anti-mouse** | 7076 | Cell Signaling Technology | Western |
| **CAMKK2** | HPA017389 | Sigma-Aldrich | Western |
| **UAP1** | HPA014659 | Sigma-Aldrich | Western |
| **OGA** | HPA036141 | Sigma-Aldrich | Western |
| **COPS3** | HPA021997 | Sigma-Aldrich | Western |
| **Mdm2** | OP46 | EMD Millipore | Western |
| **Alexa flour 594 Goat Anti Mouse** | A11020 | Invitrogen | IF |
| **Alexa flour 488 Goat Anti Rabbit** | A11070 | Invitrogen | IF |

**Supplementary Table 2**

| ChIP-qPCR primers | Forward primer | Reverse primer |
|---|---|---|
| **CAMKK2 promoter** | AGAACACTGTAGCTCACACAGGCA | GGGCACTTCCCAACCTTTCTTACT |
| **Negative AR ChIP** | AACTCCACATTTCCTAAGTGACC | CCAACCCACACCAAGTACC |
| **VPRBP promoter** | CTTGTTCTGACCTGTGTGCG | GAGGCGGAGGTCATAGTGAG |
| **Negative RL2 ChIP** | AACTCCACATTTCCTAAGTGACC | CCAACCCACACCAAGTACC |
| **PAICS/PPAT** | AGCGTACCAAGTCTTGACGG | CAGCACCCAACAGTAGCGTA |
| **p21 ChIP** | ACAGCCAGAAGCTCCAAAAA | TGCCACACACCAGTGACTTT |
| **Negative primer1 p53 ChIP** | AACTCCACATTTCCTAAGTGACC | CCAACCCACACCAAGTACC |
| **Negative primer2 p53 ChIP** | TGCCACACACCAGTGACT TT | ACAGCCAGAAGCTCCAAAAA |

**Supplementary Table 3**

| Sigma KiCqStart™ PrimerPair ID | Gene Symbol | Gene ID | RefSeq ID | Exons | Species |
|---|---|---|---|---|---|
| **H_VPRBP_1** | VPRBP | 9730 | NM_001171904 | 7-8 | Human |
| **H_CDKN1A_1** | CDKN1A | 1026 | NM_000389 | 2-3 | Human |
| **H_MGEA5_1** | MGEA5 | 10724 | NM_001142434 | 7-8 | Human |
| **H_COPS3_1** | COPS3 | 8533 | NM_001199125 | 11-12 | Human |

**Supplementary Table 4**

| Gene | Forward primer | Reverse primer |
|---|---|---|
| **CAMKK2** | TGAAGACCAGGCCCGTTTCTACTT | TGGAAGGTTTGATGTCACGGTGGA |
| **UAP1** | TCCCCACTAAAGAATGCT | AATTGCTGGAAGGCGAGA |
| **OGT** | GCAGCAGGACCAATTACCTC | GCATACGTTTCGTTGGTTCTG |
| **TP53** | GATGGTGGTACAGTCAGAGCC | TCCGAGTGGAAGGAAATTTGC |
| **RPLPO** | ATCAACGGGTACAAACGAGTC | CAGATGGATCAGCCAAGAAGG |

**Supplementary Table 5**

| siRNA | Assay ID | Catalog number | Manufacturer |
|---|---|---|---|
| **OGT si1** | s16094 | 4390824 | Thermo Fisher Scientific |
| **OGT si2** | s16095 | 4390824 | Thermo Fisher Scientific |
| **DCAF1 si1** | s18762 | 4392420 | Thermo Fisher Scientific |
| **DCAF1 si2** | s18763 | 4392420 | Thermo Fisher Scientific |
| **COPS3 si1** | s16227 | 4392420 | Thermo Fisher Scientific |
| **COPS3 si2** | s16229 | 4392420 | Thermo Fisher Scientific |
| **Silencer™ Select Negative Control No. 1 siRNA** | | 4390843 | Thermo Fisher Scientific |

**Supplementary Table 6**

| Parameter | n evaluable | negative (%) | weak (%) | moderate (%) | strong (%) | p value |
|---|---|---|---|---|---|---|
| **All cancers** | 2216 | 5.9 | 10.3 | 37.6 | 46.2 | |
| **Tumor stage** | | | | | | |
| pT2 | 1444 | 8 | 11 | 35.5 | 45.6 | <0.0001 |
| pT3a | 468 | 3.2 | 7.1 | 38.9 | 50.9 | |
| pT3b-pT4 | 277 | 0.4 | 11.6 | 46.6 | 41.5 | |
| **Gleason grade** | | | | | | |
| ≤3+3 | 994 | 9.4 | 12.6 | 36.4 | 41.6 | |
| 3+4 | 998 | 3.5 | 8.4 | 37.7 | 50.4 | |
| 3+4 Tert.5 | 1 | 0 | 100 | 0 | 0 | <0.0001 |
| 4+3 | 188 | 1.1 | 7.4 | 42 | 49.5 | |
| 4+3 Tert.5 | 4 | 0 | 25 | 50 | 25 | |
| ≥4+4 | 28 | 3.6 | 10.7 | 46.4 | 39.3 | |
| **quantitative Gleason grade** | | | | | | |
| ≤3+3 | 994 | 9.4 | 12.6 | 36.4 | 41.6 | |
| 3+4 ≤5% | 213 | 6.1 | 7 | 31.9 | 54.9 | |
| 3+4 6-10% | 241 | 4.1 | 12.4 | 34.9 | 48.5 | |
| 3+4 11-20% | 162 | 2.5 | 8 | 39.5 | 50 | |
| 3+4 21-30% | 112 | 1.8 | 7.1 | 42 | 49.1 | |
| 3+4 31-49% | 77 | 2.6 | 6.5 | 36.4 | 54.5 | 0.0001 |
| 3+4 Tert.5 | 1 | 0 | 100 | 0 | 0 | |
| 4+3 50-60% | 62 | 1.6 | 6.5 | 38.7 | 53.2 | |
| 4+3 Tert.5 | 4 | 0 | 25 | 50 | 25 | |
| 4+3 61-100% | 60 | 1.7 | 11.7 | 40 | 46.7 | |
| ≥4+4 | 6 | 0 | 16.7 | 16.7 | 66.7 | |
| **Lymph node metastasis** | | | | | | |
| N0 | 1083 | 3.9 | 9.8 | 41 | 45.3 | 0.392 |
| N+ | 65 | 1.5 | 15.4 | 41.5 | 41.5 | |
| **Preoperative PSA level (ng/ml)** | | | | | | |
| <4 | 333 | 6 | 8.1 | 36.3 | 49.5 | 0.3407 |
| 04-Oct | 1203 | 5.9 | 10.2 | 36.2 | 47.7 | |
| Oct-20 | 465 | 5.8 | 11.6 | 41.3 | 41.3 | |
| >20 | 169 | 7.1 | 10.7 | 40.2 | 42 | |
| **Surgical margin** | | | | | | |
| negative | 1707 | 6.9 | 9.8 | 38 | 45.2 | 0.001 |
| positive | 482 | 2.7 | 11.6 | 36.1 | 49.6 | |
